# The type VI secretion system governs strain maintenance in a wild mammalian gut microbiome

**DOI:** 10.1101/2025.11.29.690828

**Authors:** Beth A. Shen, Kyle L. Asfahl, Bentley Lim, Savannah K. Bertolli, Samuel S. Minot, Matthew C. Radey, Kelsi Penewit, Billy Ngo, Stephen J. Salipante, Christopher D. Johnston, S. Brook Peterson, Andrew L. Goodman, Joseph D. Mougous

## Abstract

Bacteria inhabiting the mammalian gut coexist in dense communities where contact-dependent antagonism mechanisms are widespread. The type VI secretion system (T6SS) is an interbacterial toxin delivery pathway prevalent among gut Bacteroidales, yet its function in naturally evolved microbiomes remains poorly defined. Here, we examine the role of the T6SS in *Bacteroides* within a physiologically relevant gut community derived from wild mice (the WildR microbiome). Using newly developed genetic tools and a strategy for functional replacement of strains within the WildR community, we demonstrate that the WildR isolate *B. acidifaciens* employs a T6SS to antagonize co-resident Bacteroidales. We also show that loss of T6SS function compromises the long-term maintenance of *B. acidifaciens* in the community but not its initial colonization, establishing the system as a determinant of strain persistence. The T6SS we identified resides on an integrative and conjugative element (ICE). ICE-seq, a targeted sequencing approach, reveals that the T6SS-ICE is distributed among select *Bacteroidales* and *Muribaculaceae* species in the WildR microbiome, between which it appears to be recently exchanged. We also show that transfer of the T6SS-ICE to WildR isolate *Phocaeicola vulgatus* confers transient colonization benefits in mice, but is linked to eventual population decline. Our findings demonstrate that the T6SS can stabilize the presence of specific strains within a complex, co-evolved gut microbiome, yet its value is context dependent and constrained by the ecological and physiological landscape of the host community.

## Introduction

Pathways for contact-dependent antagonism between bacteria are prevalent among species residing in the densely colonized mammalian gut (Coyne et al. 2016, Verster et al. 2017, Whitney et al. 2017, Garcia-Bayona and Comstock 2018). Prominent among these is the bacterial type VI secretion system (T6SS), a mechanism employed by Gram-negative species to deliver toxic effector proteins directly to contacting Gram-negative cells (Hood et al. 2010, Schwarz et al. 2010, Coulthurst 2019). The T6SS is widely distributed among gut resident species belonging to the Bacteroidota phylum. Over half of *Bacteroides* spp. isolated from mammalian gut samples encode T6SS genes, and Bacteroidales T6SS effector genes can be found in >60% of individual human metagenomic datasets (Verster et al. 2017, Garcia-Bayona et al. 2021). *In vitro* experiments suggest the T6SS of *Bacteroides* spp. can target a broad cross section of gut Bacteroidales, while it does not appear to act on Proteobacteria, the second most abundant group of Gram-negative organisms in the gut (Chatzidaki-Livanis et al. 2016).

A number of model murine gut microbiome studies have sought to provide insight to the role of the Bacteriodales T6SS in the gut. Studies employing gnotobiotic animals co-colonized by pairs of strains demonstrate that T6SS-mediated killing can occur in this environment, and show that targeting in laboratory co-cultures is not necessarily diagnostic of an interaction between strains in the murine intestine(Chatzidaki-Livanis et al. 2016, Wexler et al. 2016, Ross et al. 2019, Robitaille et al. 2023, Sheahan et al. 2024). Gnotobiotic experiments in which more complex mixtures of bacteria (up to 12 species) are co-inoculated and colonization of conventional mice with pairs of *Bacteroides* strains demonstrate T6SS-mediated antagonism can occur in the context of a more diverse assemblage (Hecht et al. 2016, Wexler et al. 2016, Hill et al. 2024). However, these experiments all suffer from a number of limitations. For instance, the species employed are not native to the mouse gut, and the antagonistic interactions being assessed are between strains that may have never co-inhabited a natural community. Gnotobiotic experiments additionally suffer from the lack of community complexity, while experiments employing conventional animals require antibiotic pre-treatment or large inoculums to enable strain engraftment (Hecht et al. 2016, Hill et al. 2024). As a result of these limitations, the physiological and ecological role of the T6SS in mammalian gut microbial communities remains largely unknown.

The limited insights available into the function of the T6SS in native gut microbiomes primarily derive from metagenomic analyses of T6SS gene distribution patterns. T6SS gene clusters in Bacteroidales segregate into three Genetic Architectures (GA) 1-3 based on synteny and evolutionary relatedness; however, each contain conserved structural genes and variable effector and cognate immunity genes (Coyne et al. 2016). GA1 and GA2 loci are encoded on mobile elements, and several studies show they can be horizontally exchanged among Bacteroidales strains co-resident in a microbiome, suggesting a selective benefit to obtaining T6SS effector and immunity genes shared by neighboring species (Coyne et al. 2014, Garcia-Bayona et al. 2021, Sheahan et al. 2024). Further evidence supporting selection for compatibility among the T6SS effector repertoire in co-resident species lies in the finding that effector gene diversity in individual microbiomes is low, in contrast to the high degree of effector diversity observed across metagenomes (Verster et al. 2017). Studies have also revealed that gut Bacteroidales species can encode clusters of T6SS immunity genes unlinked to cognate effectors (Ross et al. 2019). One type of these acquired interbacterial defense (AID) gene clusters, designated rAID for recombinase-associated, are widely distributed across Bacteroidales and have features consistent with active acquisition of additional immunity genes. A subset of genes encoded within AID and rAID systems selectively neutralize T6SS toxins, indicative of the selective benefit of resisting T6SS-based intoxication in the gut environment. However, other evidence suggests that under certain conditions the cost of producing a T6SS can outweigh its benefit. Studies of the prevalence of T6SS genes among *B. fragilis* strains reveal they are more common in the microbiomes of infants than in adults (Verster et al. 2017, Robitaille et al. 2023). Furthermore, longitudinal sampling of individual human gut microbiomes has revealed instances in which the *B. fragilis* T6SS acquires inactivating mutations over time(Robitaille et al. 2023). A comprehensive understanding of the physiological role of the T6SS in gut *Bacteroidales* has thus yet to emerge.

Experimental testing of the hypotheses regarding T6SS function generated from metagenomic and theoretical studies requires a tractable experimental model substantially more representative of natural gut communities than those that have been employed to date. Here, we study the function of a Bacteroidales T6SS in a natural gut community derived from wild mice and reintroduced into gnotobiotic animals (Rosshart et al. 2017). We show that this community retains a similar composition and level of diversity as the wild mouse community from which it derives, and we demonstrate we can functionally replace members of the community by adding an excess of a mutant strain of interest at the time of community introduction. Using this approach, we demonstrate that the T6SS of *B. acidifaciens* is important for maintaining this species in the community. We also demonstrate T6SS-mediated targeting between *Bacteroides* species from the same gut microbiome, and provide evidence that not all Bacteroidales in the community benefit from horizontal acquisition of the T6SS. Together, our findings show that the T6SS can be an important fitness determinant in a native gut community of naturally co-occuring organisms, and that its contribution to competitiveness in the gut varies across species.

## Results

### The WildR microbiome is stable and similar to wild mice microbiomes despite propagation

The WildR microbial community consists of a complex assemblage of organisms obtained from the pooled ileocecal contents of three wild mice and subsequently introduced to germ-free mice for laboratory propagation (Rosshart et al. 2017). This community diverges substantially from that found in lab-reared mice, with a significantly greater proportion of strains from the Bacteroidota and Pseudomonodata phyla, and a lower proportion of Bacillota and Verrucomicrobia. Additionally, the most abundant OTUs in the WildR population are consistently missing from lab mouse microbiomes. Importantly, by the F4 generation of laboratory propagation in mice, much of the diversity of the WildR was retained, and its composition was indistinguishable from that found in the originally gavaged dams (Rosshart et al. 2017).

To establish whether mice colonized by the WildR community could serve as an appropriate model to study T6SS-mediated interactions in the gut, we first sought to determine whether community composition diverged in the subsequent generations of propagation since its initial study. We obtained ileocecal contents of the F7 WildR generation, introduced these to gnotobiotic mice via oral gavage, then harvested and cryopreserved cecal contents after 14 days of *in vivo* propagation of the community (Supplemental Figure 1A). To assess the phylogenetic composition of the WildR microbiome after this propagation, we performed short-read metagenomic sequencing on DNA extracted from two stored aliquots and two fecal pellets from mice used to propagate the community, generating a total of 63 Gb of sequence data. Reads were then mapped to a mouse microbiome-specific reference database using Kraken 2 (Wood et al. 2019). This analysis revealed that, like the initial wild mouse microbiome samples and F2 generation, the WildR F7 community is enriched for Bacteroidota and depleted in Bacillota compared to that of lab reared mice (Figure 1A, Supplemental Table 1). Comparison of the relative abundance of the 100 most prevalent genera in the wild donor mice across each community indicated extensive similarities between the wild, WildR F2 and Wild F7 communities (Figure 1B, C, Supplemental Figure 1B). In contrast, lab mouse microbiomes diverge substantially from these communities, and lack 24 genera found in each wild-mouse derived microbiome (Figure 1D, E, Supplemental Figure 1C).

**Figure 1.**
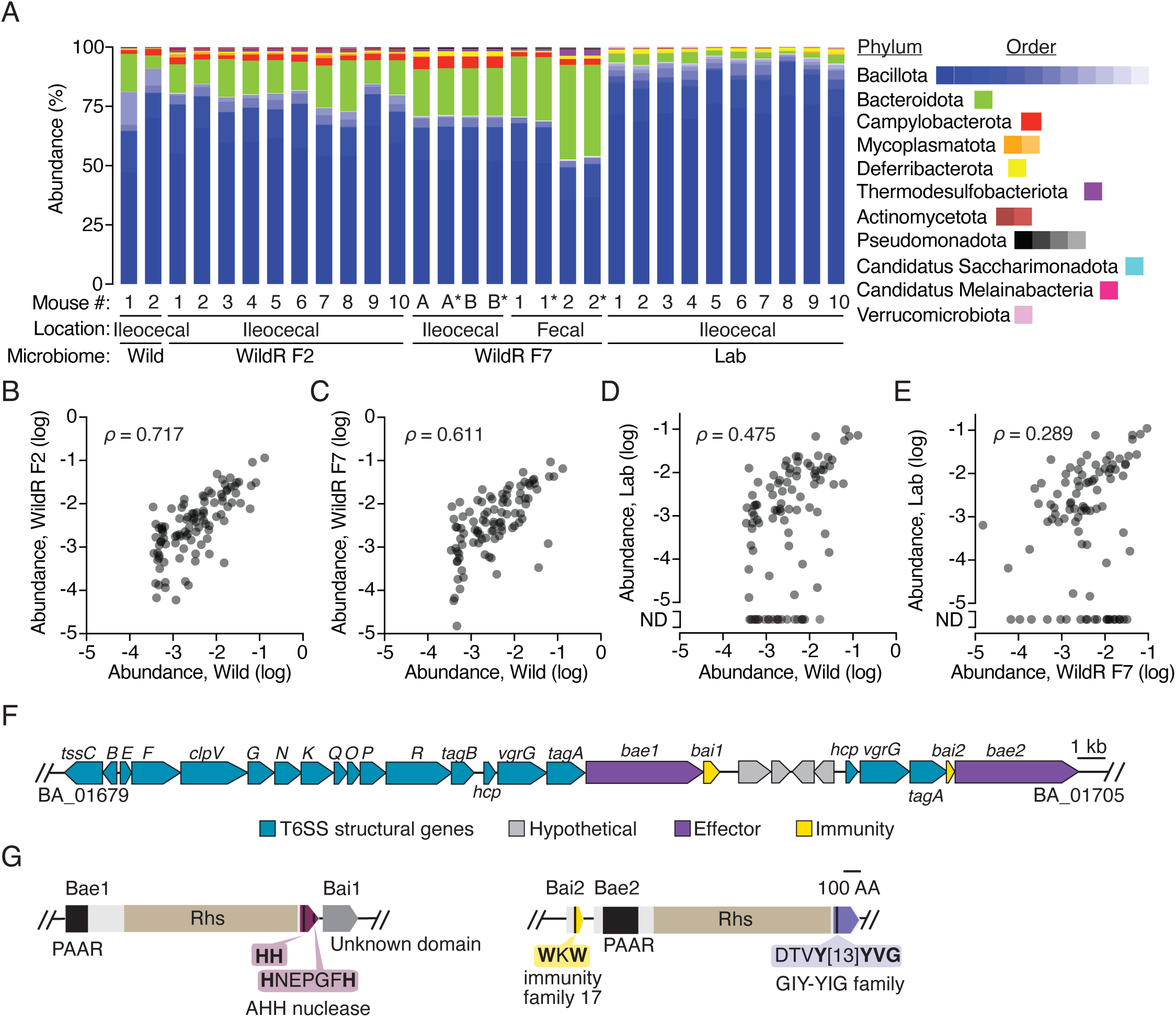
The WildR murine gut microbiome is stable over multiple generations and harbors an ICE-encoded T6SS. (A) Phylum-level taxonomic composition of the WildR or lab-derived murine gut microbiome at different generations. Data from the initial wild-caught donor mice (Wild), the WildR F2 generation and the lab mice-derived community derive from new analysis of previously published sequencing data (Rosshart et al. 2017). WildR F7 data derives from two cryopreserved samples of pooled ilocecal contents (labeled A and B) and fecal pellets from two mice (numbers 1 and 2) used to propagate the community; *indicates technical duplicate samples included in WildR F7 sequencing. Source data is presented in Supplemental Table 1. (B-E) Comparison of the abundance of the 100 genera most prevalent in wild donor mice between the indicated communities. Points are shaded to show overlapping datapoints (darker shades), and Spearman’s correlation for each comparison is indicated (ρ). ND, not detected. (F) To-scale schematic depicting the *Bacteroides acidifaciens* T6SS gene cluster. (G) Bioinformatic domain predictions for T6SS effector and immunity proteins of *B. acidifaciens*. Conserved amino acids are indicated in bold.

### Two WildR species share a mobile T6SS

Having determined that the overall community composition and diversity found in wild mouse microbiomes is preserved in the WildR F7 generation, we aimed to identify organisms in the community that encode T6SS genes. Toward this end, we used metagenomic and long-read sequencing data from WildR F7 samples to assemble high quality metagenome associated genomes (MAGs). In parallel, we obtained genome sequences for 15 of the 17 strains readily isolated from the community (Supplemental Table S6). In total, we generated nine closed and six draft genomes from isolates and 72 MAGs with an average contig N50 of 946 kb (Supplemental Table 2). Collectively, these genomes represent ∼50% of the species level diversity present in the community (Supplemental Figure 1D).

Within our collection of WildR-derived genomes, we identified three encoding T6SS genes. These include genomes from isolates of *B. acidfaciens* and *B. caecimuris* and a MAG identified as deriving from *Mucispirillum schaedleri.* The latter species is a member of the Chrysiogenota phylum (formerly Deferribacteres) for which no genetic tools are available. Thus, with the aim to functionally investigate the role of the T6SS using the WildR model, we focused on the more tractable *Bacteroides* spp. systems. We found that the T6SSs present in the WildR *B. acidfaciens* and *B. caecimuris* strains we isolated exhibit the GA1 architecture and are nearly identical across the gene cluster (Figure 1F). Like GA1 systems identified in other species, the T6SS appears to be encoded on an integrative and conjugative element (ICE) (Coyne et al. 2016). The ICE encompasses a 115 kb region, includes genes coding for DNA transfer and assorted other functions, and is identical between the genomes except for a one base-pair deletion within a predicted effector of the *B. caecimuris* GA1 cluster (Supplemental Figure 1E). This degree of similarity is consistent with recent exchange of the element between the species; previous studies of human microbiome isolates found that GA1 elements from strains co-resident in the same individual share >99% sequence identity, while those of strains isolated from different individuals share 95-98% identity(Coyne et al. 2016, Garcia-Bayona et al. 2021). Interestingly, the T6SS we found in *B. acidifaciens* and *B. caecimuris* shares only 82-83% identity with previously identified GA1 systems found in human isolates, suggesting the ICE is not being frequently exchanged between human and murine-adapted *Bacteroides* species (Supplemental Table 3).

Within the *B. acidfaciens* and *B. caecimuris* T6SS loci, we identified two candidate secreted effector genes (BA_01695 and BA_01705 in *B. acidifaciens*; Figure 1F, G). The predicted toxins contain N-terminal PAAR and Rhs domains, which are common among T6SS effectors. The C-terminal toxin domains of these proteins share no significant sequence homology or predicted structural similarity with characterized proteins. However, conserved domain analysis and comparison to structural models in the AlphaFold database indicated that both bear features of predicted nuclease toxins. BA_01695, herein renamed Bae1 (*B. acidifaciens* effector 1), contains the AHH predicted nuclease domain, while the toxin domain of BA_01705, renamed Bae2, resembles the enzymatic domain of endonucleases in the GIY-YIG family. Genes conferring immunity to T6SS system toxins are typically small open reading frames (ORFs) encoded immediately downstream of their cognate effector. Indeed, we found a putative immunity gene downstream of *bae1* (BA_01696, renamed *bai1*); a candidate immunity gene for Bae2 is located just upstream of the toxin gene (BA_01704, renamed *bai2*), and encodes a protein domain previously classified as an immunity determinant (Figure 1G). Interestingly, the single base-pair difference between the T6SSs of *B. acidifaciens* and *B. caecimuris* occurs within the *bae1* gene of *B. caecimuris*, introducing a likely inactivating frameshift (Supplemental Figure 1E). Therefore, we focused our initial functional studies of the T6SS in the WildR community on *B. acidifaciens*.

### The *B. acidifaciens* T6SS intoxicates WildR Bacteroidales

As a first step toward understanding the role played by the *B. acidifaciens* T6SS in WildR-colonized mice, we sought to determine which, if any, members of the community are susceptible to intoxication by this system *in vitro*. However, these efforts were initially confounded by our inability to genetically manipulate the WildR *B. acidifaciens* isolate using previously published approaches (Garcia-Bayona and Comstock 2019, Bencivenga-Barry et al. 2020). We hypothesized that, as in other wild isolates, restriction modification (RM) systems constitute a significant barrier to the introduction of foreign DNA in this strain (Johnston et al. 2019). To circumvent this problem, we applied the previously described RM-silencing SyngenicDNA approach (Johnston et al. 2019). Using single-molecule sequencing, we identified eight distinct methylated motifs in *B. acidifaciens* (Supplemental Table 4), representing a complex RM system barrier within this strain. Elimination of the single methylated motif found in the integrative plasmid pNUB2 enabled us to obtain transconjugants with the plasmid harboring a *B. acidifaciens* gene of interest (*tssC*), which was otherwise recalcitrant to transfer (Supplemental Figure 2A). Although conjugation is often considered relatively insensitive to RM barriers, these data indicate that the RM landscape can substantially limit conjugative plasmid transfer in wild Bacteroides (Dimitriu et al. 2024). Thus, we applied this same RM silencing approach to generate a series of improved plasmids for allelic exchange, which were used to successfully inactivate several T6SS effector, immunity and structural genes from *B. acidifaciens*.

To confirm that Bae1 and Bae2 are active T6SS effectors, we performed *in vitro* growth competition assays under contact-promoting conditions using wild-type *B. acidifaciens* and mutants sensitized to one or both of the toxins by deletion of the predicted immunity determinants. These experiments revealed that co-cultivation with the wild-type strain inhibits the growth of mutants sensitized to Bae1 or Bae2 through immunity gene inactivation (Figure 2A). A strain lacking a functional T6SS (*B. acidifaciens* Δ*tssC*) did not inhibit the effector-sensitized strains, indicating both Bae1 and Bae2 are delivered via the T6SS and have the capacity to inhibit target cell growth.

**Figure 2.**
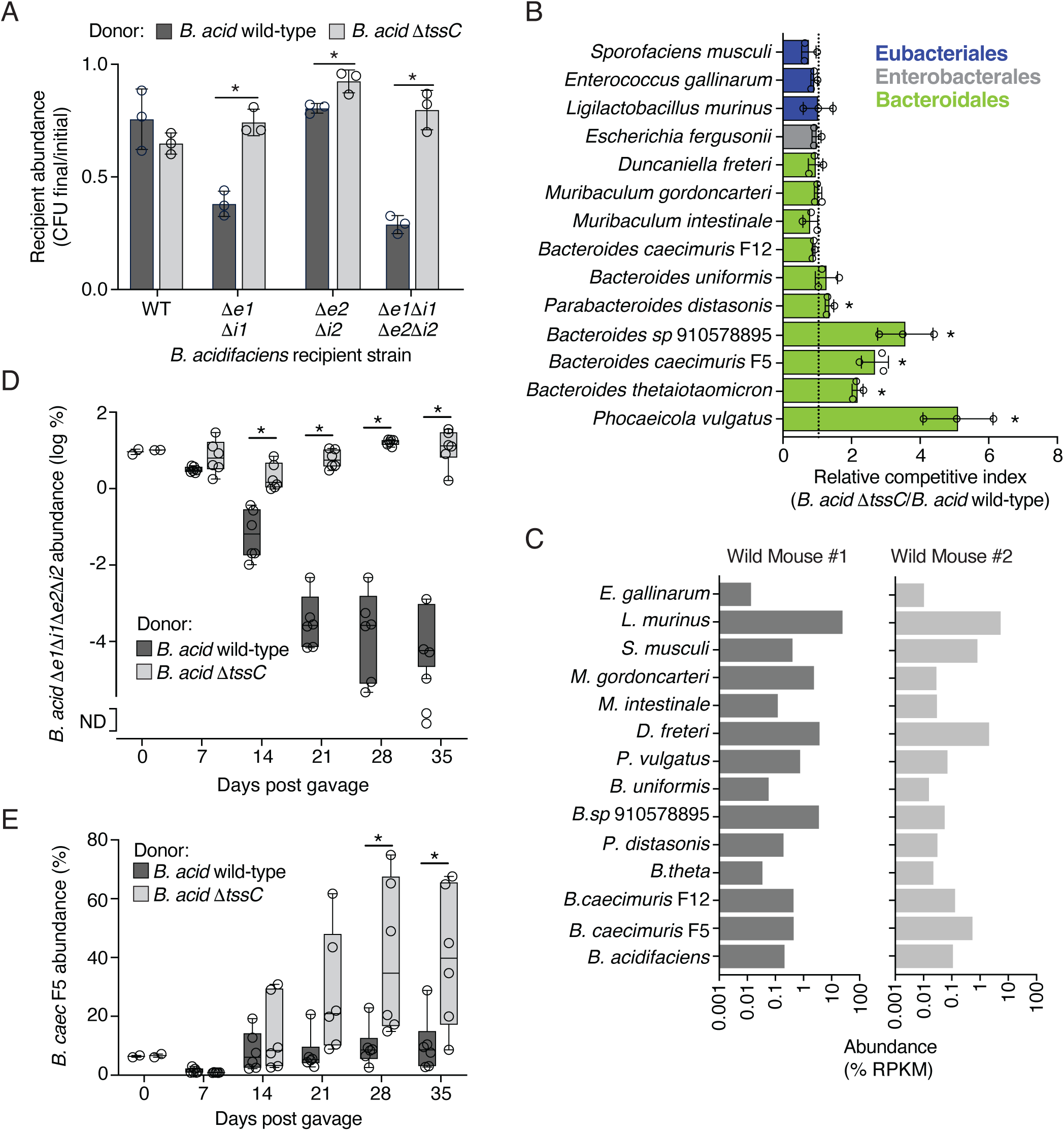
**The *B. acidifaciens* T6SS intoxicates other WildR Bacteroidales species.**(A) Normalized growth yield (final/initial CFU, mean ± SD) of the indicated strains of *B. acidifaciens* (recipients) after *in vitro* growth in competition with *B. acidifaciens* wild-type or Δ*tssC* (donors). Data are representative of at least three biological replicates. \**p* ≤ 0.05 (two-tailed t-test). CFU, colony forming units; *bae, e; bai, i*. (B) Mean (± SD) relative fitness of wild-type *B. acidifaciens* relative to a Δ*tssC* derivative during *in vitro* growth competition assays with the indicated species isolated from the WildR community. Data represent 3 biological replicates. *indicates the T6SS is significantly important for competitiveness (unpaired two-tailed t-test. *p* < 0.05). (C) Abundance (% reads per kilobase per million) of WildR species in two initial wild-caught donor mice used to establish the WildR (Rosshart et al. 2017). (D) Abundance of *B. acidifaciens* Δ*bae1* Δ*bai1* Δ*bae2* Δ*bai2* in mouse fecal samples collected following co-gavage of germ-free mice with this strain and wild-type or T6SS-inactivated (Δ*tssC*) *B. acidifaciens.* ND, not detected. \**p* < 0.05 (mixed-model ANOVA with repeated measures and Šidák’s multiple comparisons tests) (E) Relative abundance (compared to total *Bacteroides*) of *B. caecimuris* F5 following co-gavage of germ-free mice with *B. acidifaciens* wild-type or Δ*tssC*. \**p* < 0.05 (two-way ANOVA with repeated measures and Šidák’s multiple comparisons tests). For D and E, n=6 mice from two independent replicates. Boxplots represent the interquartile range and mean for each condition; whiskers represent minimum and maximum detectable values; points represent individual values.

Previous studies indicate the Bacteroidales T6SS mediates targeting of other Bacteroidales, but the exploration of targeting beyond this order has been limited to *E. coli*, which is not intoxicated (Russell et al. 2014, Chatzidaki-Livanis et al. 2016, Wexler et al. 2016). To explore the extent of T6SS-mediated intoxication in our system, we used in vitro competition assays to measure *B. acidifaciens* T6SS targeting of each of the species we isolated from the WildR community. Like the T6SS of other Bacteroidales, we found that targets of the *B. acidifaciens* system were restricted to members of the same order (Figure 2B) (Russell et al. 2014, Chatzidaki-Livanis et al. 2016, Wexler et al. 2016). Given that the fitness benefit conferred by the T6SS in these experiments was relatively modest (<10 fold), we also generated two additional T6SS inactivated mutants of *B. acidifaciens* (Δ*clpV* and Δ*tssN*). These strains exhibited a similar competitive defect against *B. caeicimuris* F5 and *Phocaeicola vulgatus* as *B. acidifaciens* Δ*tssC*, and the phenotype could be complemented by heterologous expression of the inactivated genes (Supplemental Figure 2B, C). There were also Bacteroidales for which we were unable to observe evidence of targeting by the *B. acidifaciens* T6SS. As expected, the *B. caecimuris* strain F12, which carries homologs of *bai1* and *bai2* within its T6SS gene cluster, is resistant to targeting. With the exception of *B. uniformis*, the other strains resistant to target are Bacteroidales from outside of the genus *Bacteroides*. Importantly, examination of the metagenomic data from two of the wild mice used to generate the WildR revealed that *B. acidifaciens*, and each of the candidate target species we tested were detected in both animals, indicating the interaction patterns we observe are reflective of T6SS targeting between species co-resident within a natural microbiome (Figure 2C). Together, these results demonstrate the potential for T6SS-mediated targeting between members of the same natural gut community.

Previous studies indicate that the range of T6SS-mediated targeting observed for Bacteroides strains during *in vitro* growth assays does not necessarily predict *in vivo* interactions (Wexler et al. 2016). Thus, we used co-colonization of gnotobiotic mice assays to evaluate the *B. acidifaciens* T6SS targeting capacity in the murine gut. As a first test of T6SS activity in this environment, we assessed the ability of *B. acidifaciens* wild-type or Δ*tssC* to intoxicate an immunity-deficient target strains (Δ*bae1Δbai1*Δ*bae2Δbai2*). By 14 days after introduction of the strain pairs by co-gavage, we found the immunity-deficient strain was significantly depleted in the presence of wild-type but not T6SS-inactive *B. acidifaciens,* and decline of the immunity-deficient population continued until the end of the experiment 35 days post-gavage (Figure 2D). We then used this model to evaluate *B. acidifaciens* T6SS-mediated targeting of two strains we found to be susceptible to intoxication by the system *in vitro, B. caecimuris* F5 and *P. vulgatus.* During co-colonization with wild-type *B. acidifaciens, in vivo* growth of *B. caecimuris* F5 was largely inhibited, whereas by 28 days post gavage with *B. acidifaciens* Δ*tssC*, this strain became the predominant population in fecal samples (Figure 2E, Supplemental Figure 2D). To our knowledge, these data represent the first demonstration of *in vivo* T6SS-mediated targeting between *Bacteroidales* strains derived from the same natural gut microbiome.

In contrast to the dynamics observed during *B. acidifaciens–B. caecimuris* co-colonization experiments, we found that T6SS inactivation had little impact when *B. acidifaciens* was introduced to gnotobiotic mice along with *P. vulgatus* (Supplemental Figure 2E, F). In these experiments, *P. vulgatus* quickly became the predominant colonizer regardless of the T6SS activity of *B. acidifaciens*, representing ∼80% of the total population by 7 days post gavage. This overall level of competitiveness of *P. vulgatus* is consistent with the predominance of this species among Bacteroidales in the WildR community (Supplemental Figure 2G).

### A method for modified strain introduction into the WildR

Evaluating the role of the *B. acidifaciens* T6SS in the context of WildR colonized mice, requires a method to introduce mutant strains into the community. Prior studies of strain engraftment indicate that priority effects and competitive exclusion are barriers to introducing a derivative strain to an established microbiome containing the parent or closely related strains (Lee et al. 2013, Maldonado-Gomez et al. 2016, Obadia et al. 2017, Martinez et al. 2018, Segura Munoz et al. 2022). Additionally, previously proposed methods for manipulating microbiomes *in situ* with antibiotic treatments followed by delivery of a new strain or *in situ* genetic engineering suffer from the requirement to identify antibiotics or *in vivo* amenable genetic tools that specifically target a species of interest (Jin et al. 2022, Rubin et al. 2022, Segura Munoz et al. 2022). Given that our method of introducing the WildR community to the murine gut involves oral gavage of the mixed population and its subsequent establishment, we hypothesized that we could circumvent these challenges and facilitate colonization of *in vitro* modified *B. acidifaciens* isolate by taking advantage of the natural carrying capacity of the gut microbiome (Lee et al. 2013, Hecht et al. 2016, Segura Munoz et al. 2022). Specifically, we reasoned that the addition of an excess of a *B. acidifaciens* modified strain to the WildR community during gavage would lead to displacement of the endogenous *B. acidifaciens* population while maintaining the overall natural composition of the WildR community (Figure 3A).

**Figure 3.**
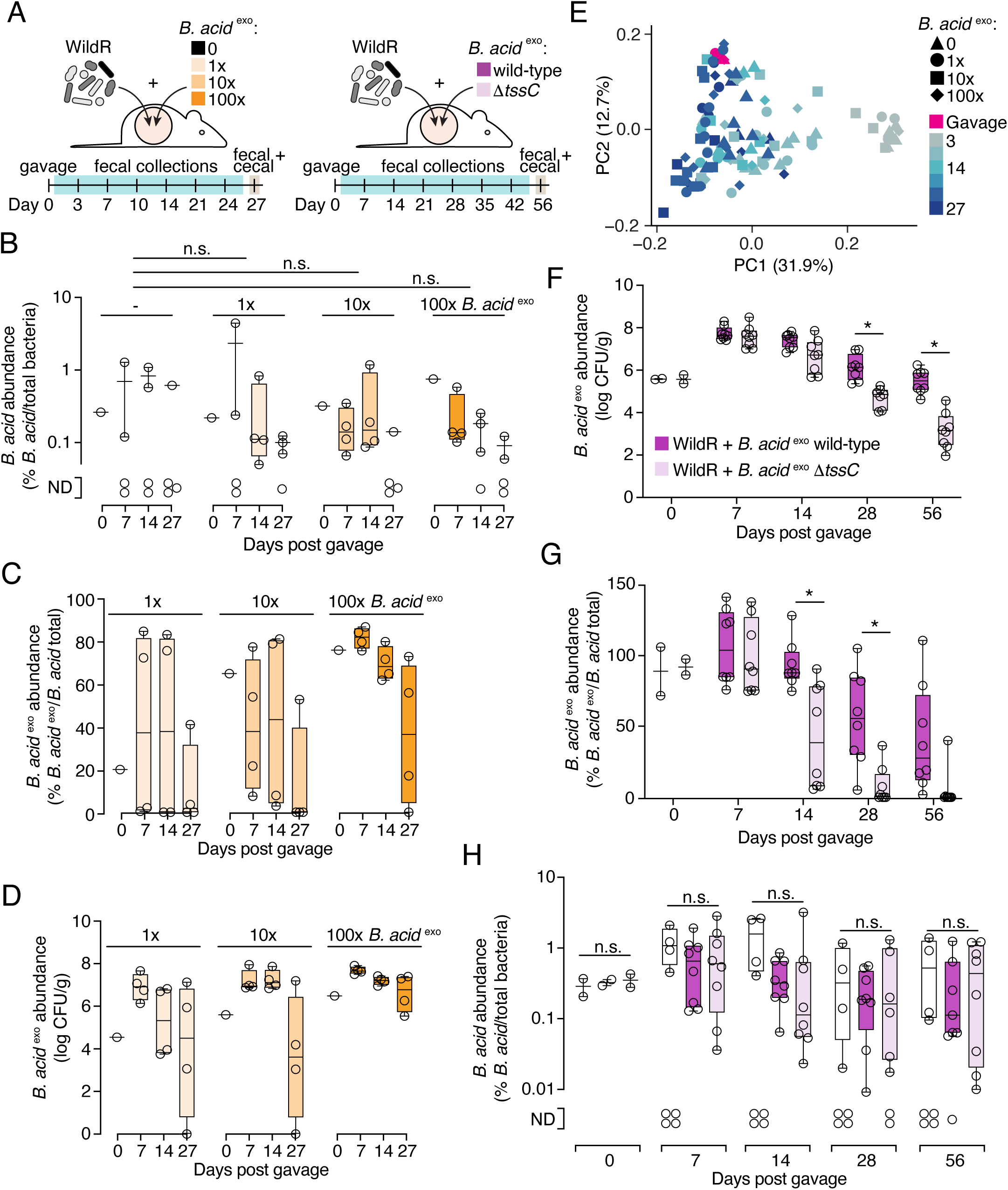
**Maintenance of *B. acidifaciens* in the WildR community is mediated by the T6SS**(A) Schematic of approach to exploit the carrying capacity of the mouse gut to promote modified strain engraftment during WildR microbiome establishment without affecting community structure. Left, design of proof-of-concept experiment to evaluate the effect of increasing the amount of *B. acidifaciens* relative to the amount of the WildR microbiome during oral gavage. Right, design of experiment to evaluate the importance of the T6SS for *B. acidifaciens* fitness in the WildR microbiome-colonized mouse gut. (B-D) Analysis of total (B), or introduced (C,D) *B. acidifaciens* populations in gavage (Day 0) or post-gavage fecal samples from germ-free mice colonized with the WildR and variable levels of *B. acid* ^exo^. B) Total abundance of *B. acidifaciens* calculated from sequencing 16S rRNA genes amplified from DNA extracted from fecal samples. Differences in *B. acidifaciens* abundance across mice gavaged with different amounts of *B. acid* ^exo^ was not significant (n=4 mice/sample, mixed-effects analysis). C) Relative abundance of *B. acid* ^exo^ in the indicated fecal samples compared to total *B. acidifaciens* as determined by qPCR. D) Abundance of *B. acid* ^exo^ in the indicated fecal samples as determined by plating for CFU on selective media. (E) Principal coordinate analysis of weighted Unifrac diversity metrics calculated from 16S rRNA gene amplicon sequencing data from feces collected from mice colonized with the WildR and variable amounts of *B. acid* ^exo^. (F-H) Quantification of *B. acid ^exo^* (F,G) or total (H) *B. acidifaciens* population in gavage (Day 0) and post-gavage fecal samples from germ-free mice colonized with the WildR community and 10-fold excess wild-type or Δ*tssC B. acid ^exo^*. n=8 mice per strain, across two biological replicates. (F) CFU quantification of *B. acid* ^exo^ and *B. acid* ^exo^ Δ*tssC*. \**p* ≤ 0.05 (two-way ANOVA with repeated measures and Šidák’s multiple comparisons test). (G) Abundance of *B. acid* ^exo^ or *B. acid* ^exo^ Δ*tssC* relative to the total *B. acidifaciens* population, as determined by qPCR. \**p* ≤ 0.05 (two-way ANOVA with repeated measures and Šidák’s multiple comparisons test). (H) Total abundance of *B. acidifaciens,* calculated from 16S rRNA gene amplicon sequencing. Samples from mice colonized by the WildR community alone (no *B. acid* ^exo^, white bars) are included for comparison. Differences in *B. acidifaciens* abundance based on *B. acid* ^exo^ genotype were not significant (mixed-model ANOVA test). For panels B-D and F-H, boxplots represent the interquartile range with indicated mean for each condition, whiskers represent maximum and minimum detectable values, and points show values from individual mice.

To test this hypothesis, we assessed the potential for *B. acidifaciens* marked with erythromycin resistance (*B. acid ^exo^*) to replace the endogenous population (*B. acid ^endo^*) when titrated into the WildR gavage mixture at different levels (Figure 3A). In cryopreserved samples of the F7 WildR community, we determined that *B. acid ^endo^*is present at ∼3×10^4^ CFU/mL (see Methods). We then added 1x, 10x, or 100x *B. acid ^exo^* relative to this population (3×10^4^-3×10^6^ CFU/mL) to aliquots of the reference stocked WildR community and inoculated mice via oral gavage. Supporting our hypothesis that endogenous *B. acidifaciens* populations represent the carrying capacity for this species in the murine gut microbiome, we found that addition of 100-fold excess *B. acidifaciens* in the gavage did not significantly increase the total relative abundance of the species in mice at 7 days post-gavage (Figure 3B). We also observed that at each titration level, we successfully replaced ∼80% of the *B. acid ^endo^* population with the marked strain; this frequency is likely an underestimate due to the detection limit of the qPCR-based method used for distinguishing the strains (Figure 3C). Higher titrations of the marked strain resulted in a trend toward lengthening its persistence in the community, although this effect was not statistically significant (Figure 3D, Supplemental Figure 3A, B). Additionally, β-diversity analyses of 16S rRNA amplicon sequences revealed that *B. acid ^exo^*addition did not impact overall community composition, regardless of the inoculum level employed (linear mixed effects model P>|z| > 0.05 for all comparisons, Figure 3E). Across all starting inocula, we observed a gradual decline in *B. acid ^exo^* populations which we hypothesize is attributable to a fitness cost to carrying the erythromycin resistance marker gene.

### Long-term maintenance of *B. acidifaciens* in the WildR community requires the T6SS

With a method in hand for introducing a genetically modified strain into the WildR community, we applied this approach toward dissecting the ecological role of the *B. acidifaciens* T6SS. Our experiment was designed to test two different possible roles that have been proposed for the T6SS of *Bacteroides* in the gut microbiome: *i*) facilitating niche establishment during initial colonization and *ii*) population maintenance once stable colonization is achieved (Hecht et al. 2016, Verster et al. 2017, Robitaille et al. 2023). Germ-free mice were co-gavaged with either *B. acid ^exo^* or *B. acid ^exo^* Δ*tssC* and the WildR community (Figure 3A). *B. acid ^exo^* populations in fecal samples were then monitored over time by plating on selective media, and the proportion of the total *B. acidifaciens* population represented by *B. acid ^exo^* was monitored by qPCR. We found that through 14 days post gavage, population levels of *B. acid ^exo^*and *B. acid ^exo^* Δ*tssC* were indistinguishable (Figure 3F, Supplemental Figure 3C). However, by day 28, *B. acid ^exo^* Δ*tssC* was significantly depleted from the community compared to the marked wild-type parent strain. By 56 days post gavage, we observed a nearly 100-fold difference in the abundance of the two strains in both fecal and cecal content samples (Supplemental Figure 3D). To confirm that inactivation of the T6SS is responsible for the depletion of *B. acid ^exo^* Δ*tssC*, and not a secondary site mutation, we sequenced the genomes of the strains employed in the experiment. *B. acid ^exo^* Δ*tssC* used in the first replicate of the experiment carried a single synonymous mutation (in *cdr*) and in the second replicate the strain bore no mutations. Of note, the marked wild-type *B. acid ^exo^* population also declined over time relative to endogenous *B. acidifaciens* levels, while the overall abundance of *B. acidifaciens* in the community remained similar throughout the experiment (Figure 3G, H); we hypothesize that the decline of *B. acid ^exo^* derives from a fitness cost associated with carrying the erythromycin resistance marker. However, this decline was modest compared to the decrease in abundance of *B. acid ^exo^* Δ*tssC*.

To determine which organisms could be the targets of the *B. acidifaciens* T6SS, we performed 16S rRNA amplicon sequencing on fecal samples from mice colonized by the two *B. acid ^exo^* strains and parallel control mice colonized with the WildR community alone. This analysis revealed no significant differences between the three communities at any of the time points sampled (Supplemental Figure 3E). One potential explanation for this finding is *B. acidifaciens* is present at significantly lower abundance in the community than the organism(s) it targets. In this scenario, T6SS-targeting could be required for *B. acidifaciens* population survival without causing a sufficiently large decrease in the population of the target to be detectable by the relatively low-resolution measurement method we employed. Consistent with this potential explanation for the lack of overall community shift upon inactivation of the *B. acidifaciens* T6SS, we find that this species is present at 5 to 10-fold lower abundance than five other Bacteroidales species in WildR F7 samples (Supplemental Figure 2G). Together, our results suggest that the T6SS is required for *B. acidifaciens* population maintenance in the gut, but dispensable for initial colonization.

### Distribution of a mobile T6SS in the WildR community

Given the fitness benefit of the T6SS we observed for *B. acidifaciens* during stable colonization of the mouse gut, and previous observations that T6SS-containing ICE can be present in multiple species in a single microbiome (Coyne et al. 2014, Zhang et al. 2024), we sought to better understand the distribution of the system in the WildR community. To avoid both the biases inherent in screening for the element in cultured isolates and the challenges of assigning repetitive sequences from metagenomic datasets to MAGs, we developed a targeted sequencing-based approach for identifying ICE-carrying strains in DNA extracted directly from WildR-colonized mouse fecal samples. Our method, which we named ICE-seq, is based on techniques used for identifying transposon insertion locations (Gallagher 2019). In short, we designed primers to specifically amplify and sequence outward from the 5’ and 3’ ends of the ICE to capture the genomic context of the insertion (e.g., ICE junctions; Supplemental Figure 4A). After amplification and sequencing of ICE junctions from fecal pellets collected from 4 WildR-colonized mice, reads were mapped to a large, dereplicated genome catalog containing the WildR MAGs generated in this study and the comprehensive mouse microbiota genome (CMMG) catalog to identify ICE-containing strains (Gallagher 2019, Kieser et al. 2022). We assigned a positive identification of an ICE-containing strain based on the presence of reads mapping from either end of the ICE that diverge from a single position on a contig.

Supporting the validity of our approach, we detected ICE insertion within *B. acidifaciens* and *B. caecimuris* F12 (Figure 4A). For these and other species, detection of ICE and the proportion of ICE-seq reads each represented varied between mice. For *B. acidifaciens* and *B. caecimuris* F12, this appears to reflect their abundance in individual mice; however, we posit that exchange of the ICE on a rapid timescale could explain the additional variability observed. We also found three additional strains harboring the ICE: *Muribaculaceae* CAG-485 sp002362485*, Bacteroides* spMGG23559, and *Phocaeicola sartorii*. In *Muribaculaceae* CAG-485 sp002362485, we identified four insertion locations for the ICE, suggesting there are multiple populations of this species harboring the element within the community. To our knowledge, T6SS-encoding ICE have not previously been detected in a Muribaculaceae family member. Strikingly, we did not find evidence of ICE insertion in many of the more abundant Bacteroidales species in the community, including *P. vulgatus, Bacteroides* sp. 910578895 and *Parabacteriodes* sp. 910577325. These findings suggest that fitness benefits conferred by acquiring the ICE could vary across species.

**Figure 4.**
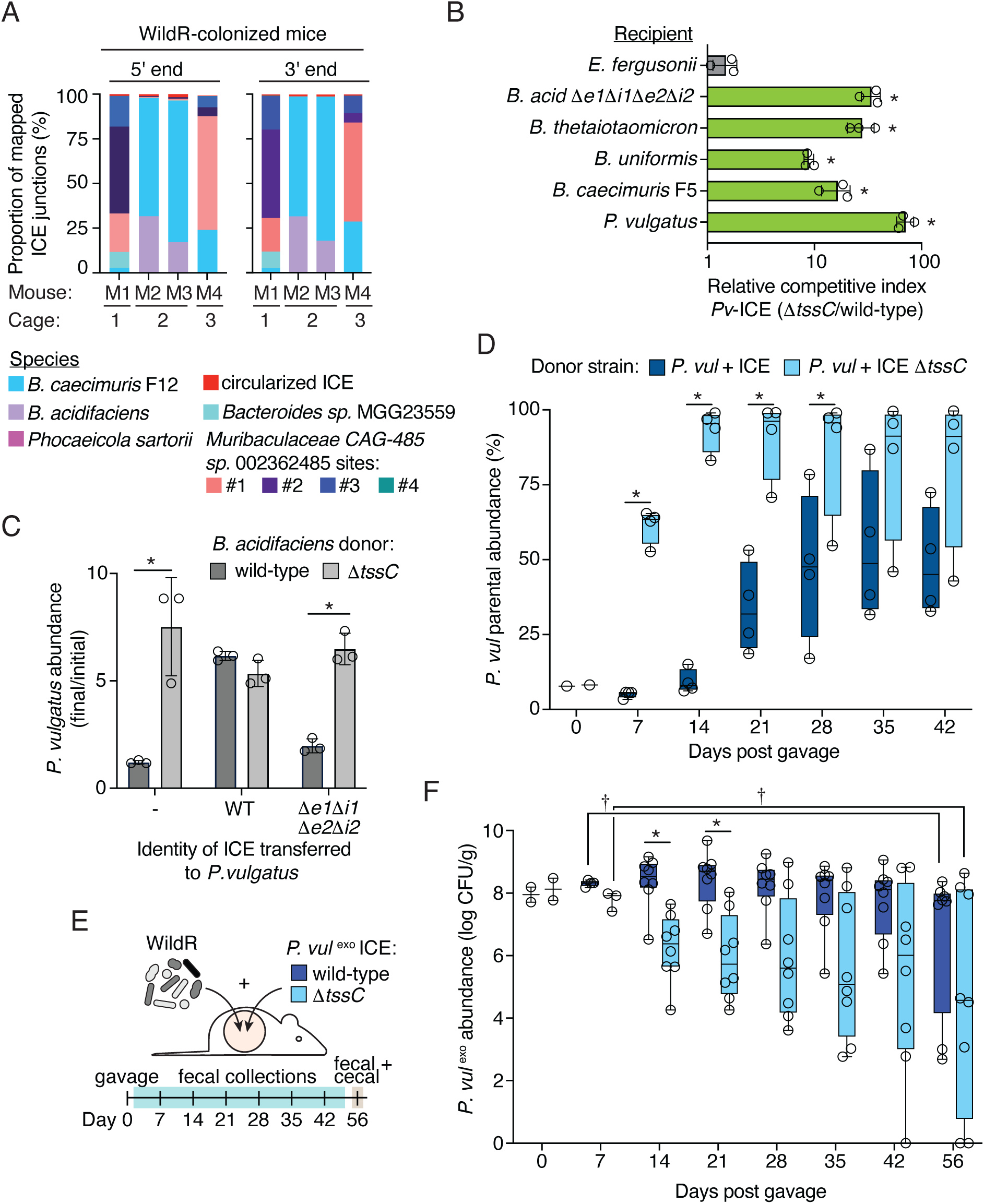
*P. vulgatus* gains a limited fitness benefit from the T6SS-encoding ICE in WildR-colonized mice. (A) Frequency of mapped ICE junctions deriving from the indicated species as determined by 5′ or 3′ ICE-Seq analysis of DNA extracted from fecal samples collected either 7 or 14 days post-gavage of the WildR into germ-free mice (n=4). Mice housed in separate or shared cages are indicated. (B) Relative competitive index from *in vitro* growth competitions between *P. vulgatus* ICE or ICE Δ*tssC* and recipient strains isolated from the WildR microbiome (Pseudomonadata, grey; Bacteroidota, green). The mean ± SD from three biological replicates is shown. Asterisks indicate recipient species for which the difference in competitive index was statistically significant between donor strains (p<0.05, unpaired two-tailed t-test). (C) Recovery of *P. vulgatus* strains containing the indicated versions of the ICE after *in vitro* growth with *B. acidifaciens*. The mean ± SD from one biological replicate and its associated technical replicates are shown, which represent results from at least three biological replicates. \**p* ≤ 0.05 (two-tailed t-test). *bae, e; bai, i*. (D) Relative abundance of wild-type *P. vulgatus* in feces collected from mice (n=4) co-colonized with *P. vulgatus* ICE or *P. vulgatus* ICE Δ*tssC,* as determined by qPCR assays with primers specific for *P. vulgatus* ^exo^ or the parent endogenous strain. Boxplots represent the interquartile range with indicated mean for each condition, whiskers represent minimum and maximum values, points show values from individual mice. \**p* ≤ 0.05 (two-way ANOVA with repeated measures test and Šidák’s multiple comparisons test). (E) Schematic of experimental design to test the fitness of *P. vu*l ^exo^ + T6SS-ICE in the WildR community. (F) Recovery of *P. vul* ^exo^ + ICE (dark blue) or *P. vul* ^exo^ + ICE Δ*tssC* (light blue) from post-gavage fecal samples of mice co-colonized with the WildR community as depicted in (E). CFUs were determined by plating on selective media (erm). Boxplots represent the interquartile range with indicated mean for each condition, whiskers represent minimum and maximum values, points show values from individual mice. Data in panel F is from two biological replicates with 4 mice per group per replicate (n=8). \**p* < 0.05 (two-way ANOVA with repeated measures and Šidák’s multiple comparisons test to compare *P.vulgatus* ICE and ICE Δ*tssC* populations), † *p* < 0.05 (unpaired t-test to compare day 7 and day 56 samples).

### Acquisition of a T6SS-encoding ICE transiently benefits *P. vulgatus*

Our finding that the ICE harboring a GA1 T6SS is shared by multiple Bacteroidales species in the WildR community suggests that it can be exchanged among community members. However, we also found that many WildR Bacteroidales species do not contain the element, suggesting the fitness benefit it confers may be species-specific, or that barriers exist that prevent its transmission to certain strains. To gain insight into the mechanisms governing ICE distribution, we focused on *P. vulgatus,* a prominent member of the WildR community which our ICE-seq analysis indicates does not naturally harbor the element (Figure 4A). Using *in vitro* conjugation assays with *B. acidfaciens* in which we introduced a chloramphenicol resistance cassette to the ICE, we were able to obtain integrants of the wild-type ICE and mutant derivatives in *P. vulgatus* (Supplemental Figure 4B). Sequencing revealed the ICE inserts at multiple locations in this strain characterized by a degenerate, AT-rich sequence (Supplemental Figure 4C). These findings suggest that barriers to horizontal gene transfer are unlikely to explain the lack of acquisition of the element by *P. vulgatus* in the context of the WildR.

Having obtained *P. vulgatus* strains carrying the ICE, we next asked whether the transferred T6SS is functional in this context. Surprisingly, we found that *P. vulgatus* carrying the element is able to use the ICE-encoded T6SS to robustly antagonize a range of Bacteroidales strains isolated from the WildR, including the parent strain lacking the ICE (Figure 4B). We also found that acquisition of the ICE by *P. vulgatus* enabled this strain to resist antagonism by *B. acidifaciens* as a result of acquiring the immunity genes *bai1* and *bai2* (Figure 4C).

While our *in vitro* experiments suggest multiple potential fitness benefits for *P. vulgatus* of acquiring the ICE, the fact that we fail to detect the element in this species in WildR-colonized mice suggests it incurs a fitness cost under *in situ* conditions not captured by our *in vitro* assays. To evaluate this possibility, we co-gavaged germ-free mice with wild-type *P. vulgatus* and derivatives marked with an erythromycin resistance cassette integrated at a neutral site and carrying the intact ICE, or a version of the ICE in which the T6SS is inactivated (ICE Δ*tssC*). Strikingly, we found that despite their 10-fold excess in the gavage sample, each ICE-containing strain was eventually supplanted by the wild-type parent (Figure 4D). Replacement occurred more rapidly when the T6SS on the ICE was inactivated, indicating that T6SS mediated targeting of the parent strain likely occurred, but was insufficient to compensate for the competitive disadvantage incurred from acquisition of the element.

We next examined the persistence dynamics of *P. vulgatus* of carrying the ICE in the context of the WildR-colonized mouse intestine. Employing the same carrying capacity-mediated approach for strain engraft we developed with *B. acidifaciens*, we colonized mice with the WildR community and *P. vulgatus* strains carrying the wild-type or Δ*tssC* ICE (Figure 4E). Up to three weeks post-gavage, *P. vulgatus* carrying the wild-type ICE colonized to higher levels than *P. vulgatus* ICE Δ*tssC,* and dominated endogenous *P. vulgatus* populations (Figure 4F, Supplemental Fig 4D). This result suggests that T6SS activity provides an advantage to *P. vulgatus* early in the establishment of the gut community. However, by the conclusion of the experiment, the abundance of *P. vulgatus* ICE declined significantly, with a concomitant rise in endogenous *P. vulgatus* populations (Figure 4F, Supplemental Fig 4D). This contrasts with our findings using germ-free mice co-colonized with endogenous *P. vulgatus* and *P. vulgatus* ICE, in which we observed a cost to carrying the ICE at early time points (Figure 4D). We speculate that in the context of the WildR, the competition from the increased diversity of species exerts greater selection for antagonistic capacity. The extent of T6SS-mediated antagonism by *P. vulgatus* carrying the ICE is supported by analysis of the overall community structure. In fecal samples collected 14-21 days post gavage, when *P. vulgatus* ICE and ICE Δ*tssC* abundance differs, we observe a significant degree of variation between the two communities (Supplemental Figure 4E). At later timepoints, the communities of mice colonized by the two strains were less distinct, although they remained significantly divergent in one replicate.

The transient nature of the fitness benefit conferred to *P. vulgatus* by T6SS activity in WildR-colonized mice contrasts with the long-term benefit to T6SS activity we observed for *B. acidifaciens* (Fig 3E). Given that ICE acquisition by *P. vulgatus* would likely occur after community establishment – a state in which our data suggest the element does not provide a lasting benefit to *P. vulgatus –* these results offer an explanation for the lack of ICE in this species. Interestingly, we found that while *P. vulgatus* ICE Δ*tssC* was initially less able to colonize mice in the context of the WildR community than *P. vulgatus* carrying an active T6SS, this strain persisted at similar population levels in a subset of mice throughout the duration of the experiment (Figure 4F). This result was unexpected, given that *P. vulgatus* ICE Δ*tssC* was rapidly displaced by the endogenous *P. vulgatus* strain in pairwise gnotobiotic mouse colonization experiments (Figure 4D). This finding suggests that in the context of the WildR, functions encoded on the ICE beyond toxin delivery by the T6SS provide a benefit to *P. vulgatus*. Of note, *P. vulgatus* ICE Δ*tssC* carries the intact immunity genes and therefore retains the capacity to resist intoxication by Bae1 and Bae2 (Figure 4C); these may contribute to the fitness benefit of the element in the presence of the WildR, where these toxins could be delivered by other T6SS-ICE encoding strains.

## Discussion

In the stable mammalian gut microbiome, where individual strains can persist throughout the lifetime of an individual, the role of interbacterial antagonism pathways is not well understood. Modeling-based studies suggest that antagonistic relationships between community members can be stabilizing to the community as a whole, but how these pathways are employed in an established community and the degree to which active targeting occurs between community members in this setting is unexplored (Coyte et al. 2015, Coyte and Rakoff-Nahoum 2019). Here, we show that an active T6SS is required for persistence of the mouse commensal *B. acidifaciens* in a natural murine gut community. To our knowledge, our results represent the first evidence of T6SS-mediated targeting between naturally co-resident gut species. The species targeted by the *B. acidifaciens* T6SS *in vivo* could not be confidently ascertained by the approach we employed, likely because this strain represents a small fraction of the community and its antagonistic activity did not detectably affect other populations. However, *in vitro* experiments demonstrate that other Bacteroidales species from the same community are effectively targeted by the T6SS of this organism. We postulate that *B. acidifaciens* relies on T6SS-mediated antagonism to prevent displacement by competing *Bacteroides.* This model for the role of the T6SS in gut Bacteroidales is consistent with metagenomic analyses showing that in adult human gut microbiomes, *B. fragilis* is most likely to carry the T6SS if the overall prevalence of *Bacteroides* species is high (Verster et al. 2017, Robitaille et al. 2023). The lytic nature of T6SS-mediated killing could also contribute to the adaptive benefit of the system for Bacteroidales species by releasing nutrients from targeted organisms (Borgeaud et al. 2015, Stubbusch et al. 2025).

Under our experimental conditions, the T6SS of *B. acidifaciens* appears to be dispensable for initial colonization of the murine gut. This result was unexpected, as the T6SS is more prevalent in human infant gut microbiomes than in adults, suggesting that T6SS-mediated competition between *Bacteroides* is important during microbiome establishment (Verster et al. 2017, Robitaille et al. 2023). The colonization dynamics associated with WildR introduction to adult animals differ from the natural process of gut microbiome development. Modifications to our carrying capacity-driven approach to strain engraftment will be needed to establish a model that more faithfully replicates the early stages of natural gut microbiome establishment. Additionally, our experiments did not address the role of the T6SS in resilience following disturbance, nor in preventing strain invasion, both of which are questions that should be amenable to future studies employing the WildR model.

A feature shared by the widely distributed GA1 and GA2 T6SSs is their placement within an ICE (Coyne et al. 2016). Previous studies show these can be exchanged between species within the gut of an individual human or mouse, suggesting selection for compatibility among the effector repertoires they encode (Coyne et al. 2014, Garcia-Bayona et al. 2021, Sheahan et al. 2024). However, the factors governing the extent to which the ICE-encoded T6SSs spread among co-resident *Bacteroides* species are not well understood. We show using ICE-seq that an ICE-encoded GA1 T6SS is present in a minority of *Bacteroides* species within the gut of a natural mouse community. The ICE present in two species, *B. acidifaciens* and *B. caecimuris,* shares >99% sequence identity across the element, indicated it was recently exchanged between them. Interestingly, we find that T6SS activity contributes to *B. acidifaciens* competitiveness towards *B. caecimuris* during co-colonization of gnotobiotic mice, suggesting these species may occupy a similar niche, and thus be in direct competition in their native habitat. Accordingly, ICE acquisition by one of these species could be expected to exert strong selective pressure for acquisition and retention of the element in the second species, allowing it to resist intoxication by its competitor.

In contrast, we find that *P. vulgatus* has not acquired the T6SS-encoding ICE in the WildR gut community. We show that the T6SS is active in this species, and it can promote initial gut colonization by *P. vulgatus*. However, during stable gut colonization, the fitness cost of harboring the element appears to outweigh its benefit. One potential explanation for these results is that *P. vulgatus* relies on mechanisms other than T6SS-mediated antagonism to maintain its niche in the gut. An alternative possibility is that the fitness cost incurred by T6SS expression in *P. vulgatus* is higher than in species that maintain the element, such as *B. caecimuris* and *B. acidifaciens*. In support of this, we observed that T6SS activity provided up to a 100-fold competitive advantage over two days to *P. vulgatus* during growth in competition with WildR Bacteroidales isolates, whereas in *B. acidifaciens,* the degree of T6SS-dependent competitiveness was considerably more modest (up to five-fold). This difference suggests that *P. vulgatus* may lack a regulator that modulates the activity of the T6SS in *B. acidifaciens,* thus increasing the cost of producing the system in this species. This observation is in-line with a proposal by Comstock and colleagues that dysregulation is a factor limiting GA1 and GA2 mobility within communities (Garcia-Bayona et al. 2021). Whether other factors, such as the functions encoded by non-T6SS genes on the ICE, contribute to the differences in the fitness costs and benefits conferred by ICE acquisition across species remains to be determined.

Our finding that the T6SS of a gut *Bacteroides* species is important for stable colonization of its native habitat expands the diversity of demonstrated benefits to T6SSs in natural environments. In both the midgut of honeybees and the light organ of bobtail squid, the T6SS appears to be primarily important for intraspecies competition (Speare et al. 2018, Motta et al. 2024). Strains of the squid symbiont *Vibrio fischeri* encoding a T6SS exclude T6SS-deficient competitors from the light organ crypts they colonize (Speare et al. 2018). Similarly, T6SS-encoding honeybee gut resident *Snodgrassella alvi* excludes T6SS-deficient strains from bee gut colonization (Motta et al. 2024). In contrast, the T6SS of *Pseudomonas* species appears to be important for colonization of soil and the plant roots, and one study found that T6SS activity of *P. protegens* affected overall community composition in the tomato rhizosphere (Duran et al. 2021, Vazquez-Arias et al. 2025). Additional studies across more habitats and species will serve to build a complete picture of the ecological and evolutionary factors that have led to the dissemination of the T6SS across such a wide diversity of species.

A key challenge facing the field of microbiome research is how to investigate gut microbe functions of interest in a physiologically relevant context that recapitulates the complexity of native gut communities. In this study, we have developed a number of resources and methods to facilitate the use of WildR-colonized mice as a model microbiome composed of co-evolved species native to the murine gut environment. We have isolated 16 species from the community, and obtained genomes representing 55% of the phylogenetic diversity of the community. Our finding that the carrying capacity of the WildR community can be exploited to facilitate functional replacement of endogenous strains with modified derivatives is a generalizable method that can be used to study the role of the T6SS in other organisms, such as *Mucispirillum schaedleri,* or to address unrelated questions pertaining to other aspects of gut microbiome biology. These resources and methods should further encourage adoption of the WildR community as a valuable tool for advancing mechanistic gut microbiome research.

## Acknowledgements

We thank Ben Ross and members of the Mougous lab for helpful discussions, Stephan Rosshart for providing access to individual wild mouse microbiome data, Natasha Bencivenga-Barry for assistance with preparation of WildR cryopreserved stocks, the Microbial Interactions & Microbiome Center for providing laboratory space and equipment to support animal studies, and the UW Gnotobiotic Animal Core for assistance with gnotobiotic animal work including experimental design, breeding, and technical expertise. This work was supported by NIH grant R35GM118159 (to A.L.G) and NIH grant R01 DE027850 (to C.D.J). B.A.S. was supported by a Ruth R. Kirschstein National Research Service Award (F32 AI164853). J.D.M is a Howard Hughes Medical Institute Investigator and held the Lynn and Michael Garvey Chair in Gastroenterology at the University of Washington. J.D.M. currently holds the John F. Enders Professorship at Yale University.

## Methods

### Strains and culture conditions

Strains employed in this study are listed in Supplemental Table 5. *E. coli* strains employed include *E. coli* EC100D pir+ (for cloning), and *E. coli* S17-1 λpir and *E. coli* S17-1 λpir + RK231 (for conjugation into Bacteroidales sp.). *E. coli* was grown aerobically at 37 °C in LB. Other bacterial strains employed in this study were isolated from the WildR community as described below. Isolates and their culture conditions are listed in Supplemental Tables 5 and 6. A hard-sided anaerobic workstation (Don Whitley Scientific) containing 10% CO_2_, 10% H_2_, and 80% N_2_ was used for all anaerobic microbiology procedures. If required, antibiotics were supplemented at the following concentrations: 25 µg erythromycin (erm)/mL, 200 µg gentamicin (gent)/mL, 25 µg chloramphenicol (cm)/mL, and 100 ng anhydrotetracycline (aTc)/mL.

### Generation of WildR reference stocks

Cryoperserved stocks of the WildR microbiome were prepared as previously described with modifications as described below (Rosshart et al. 2017). All procedures were approved by the Yale University Institutional Animal Care and Use Committee (IACUC). Male and female 6-8 week old mice were housed in as singles or pairs in flexible plastic gnotobiotic isolators with at 12-hr light/dark cycle. Mice were a provided standard chow (5K67 LabDiet; Purina) and water ad libitum. Frozen ileo-cecal contents from mice colonized with the F6 generation of the WildR microbiome were obtained from Taconic Biosciences (https://www.taconic.com/services/microbiome/wild-mouse-microbiome). To propagate this community, six germ-free C57BL6 mice were inoculated with the WildR microbiome by oral gavage (100 µL/mouse) on two consecutive days. Fecal samples were collected 14 days after gavage. After 14 days post gavage, fecal samples were collected for downstream characterization, mice were euthanized and transferred to an anaerobic chamber. The contents of the ileum, cecum, and colon (“gut”) were collected and pooled from each mouse. Pooled gut contents were diluted in pre-reduced, sterile PBS-C (PBS pH 7.4 with 1% cysteine) at 15 mL/g gut contents before passing through a 70 µm cell strainer. Material was then diluted in an equal volume of storage buffer (PBS-C with 40% glycerol final concentration), aliquoted into Wheaton vials to maintain an anerobic environment, and stored at-80 °C.

### Metagenomic sequencing of the WildR community

For shotgun sequencing on the Illumina platform, genomic DNA was extracted from two aliquots of the propagated, cryopreserved WildR stock (∼250 mg of pelleted cells) and fecal pellets (∼50 mg) from two of the six mice inoculated with the Taconic WildR material using the MoBio PowerFecal DNA Isolation kit (Catalog #12830-50, MoBio Laboratories, California, USA). Sequencing libraries were prepared using an in-house protocol (Salipante et al. 2014) and sequenced on the Illumina NextSeq using the NextSeq 500/550 Highoutput kit v2.5 300 cycles (Cat # 20024908). Samples were processed in technical duplicates to correct for PCR errors. Raw reads have been deposited with links to BioProject PRJNA1367654 in the NCBI database.

For long-read sequencing (Oxford Nanopore MinION), high-molecular weight (HMW) gDNA was extracted from ∼500 µL of the WildR cryopreserved stock using the Qiagen Genomic-tip 20/G (Cat#13323) following the protocol for bacterial DNA extraction, except 5 µL of Qiagen lytic enzyme (catalog #158928) and 10 µL of MetaPolyzme (Millipore Sigma MAC4L-5MG resuspended in 750 µL of water) were added instead of lysosome for more efficient enzymatic lysis of diverse bacteria in the community (Maghini et al. 2021). HMW DNA was then cleaned and concentrated with an isopropanol precipitation as described and eluted in 100 µL of water as previously described (Maghini et al. 2021). DNA was incubated at 55 °C for 1 hr before placing at 4 °C for 3 days to increase resuspension of HMW gDNA fragments. Quality and quantity of HMW gDNA were assessed by NanoDrop UV-Vis spectroscopy and Qubit dsDNA HS fluorimetry assay (Cat# Q33231), respectively, and the presence of fragments >10 kb was confirmed by agarose gel electrophoresis. Libraries for sequencing using the Oxford Nanopore MinION were prepared using the Oxford Nanopore SQK-LSK110 kit and 902 ng of WildR HMW gDNA as the starting material. The sequencing library was loaded onto a R9.4.1 SpotON flow cell and run for 74 hrs on the Mk1C until ∼50 pores remained. The run resulted in 22 Gb of sequencing data from 8.6 million reads with an N50 of 5.3 kb. Raw reads have been deposited with links to BioProject PRJNA1367654 in the NCBI database.

### Taxonomic classification of WildR-derived metagenomic sequences

Metagenomic reads, generated from Illumina sequencing of either WildR reference stocks or fecal samples from mice 14 days post gavage with cecal contents from Taconic, were cleaned to remove adapter sequences using Trimmomatic (v0.39) (Bolger et al. 2014) and any reads mapping to the mouse (*Mus musculus*) genome (RefSeq GCF_000001635.27) using bowtie2 (Langmead and Salzberg 2012). Taxonomic classification of each read was performed using a publicly available custom Kraken2-Bracken Snakemake pipeline based on the comprehensive mouse microbiota genome catalog (CMMG; (Lu et al. 2017, Wood et al. 2019, Kieser et al. 2022) https://github.com/SilasK/Krak). To ensure high fidelity alignment, the pipeline was modified to include a confidence of 0.51 in the Kraken2 command, and a threshold of 1000 reads per sample was applied to the Bracken analysis. The same classification pipeline was applied to metagenomic sequencing data from other WildR generations and lab mice previously published by Rosshart and colleagues (Rosshart et al. 2017). Classifications were condensed at the order and genus levels for downstream analyses.

### Metagenome assembly

WildR metagenome-assembled genomes (MAGs) were generated from Illumina and MinION reads using the MUFFIN 1.0.6 pipeline ((Van Damme et al. 2021); https://github.com/RVanDamme/MUFFIN). Briefly, short and long reads were filtered for high-quality before performing hybrid assembly with MetaFlye (version 2.7; https://github.com/mikolmogorov/Flye). Instead of the default polishing step using Pilon, we modified the MUFFIN pipeline to reduce error by using Polypolish for short read polishing of our assemblies ((Wick and Holt 2022); https://github.com/rrwick/Polypolish). Binning was performed with Concot (version 1.1.0) (Alneberg et al. 2014), Metabat2 (version 2.13) (Kang et al. 2019), and Maxbin2 (version 2.2.7, https://sourceforge.net/projects/maxbin2/), and refined by metawrap (version 1.2.2) (Uritskiy et al. 2018). Hybrid reassembly was then performed with Unicycler (version 0.4.7) (Wick et al. 2017). Bin quality was determined using CheckM (version 1.0.13) (Parks et al. 2015) and taxonomic classification was performed with GTDB-tk (version 2.1.0) (Chaumeil et al. 2019). Finally, all genomes were annotated with Prokka (version 1.12) (Seemann 2014). The assembly details and quality for each MAG are listed in Supplemental Table 2. MAG sequences are available at https://doi.org/10.5281/zenodo.17716312.

### WildR isolate collection and identification

Media used for cultivation of isolates from the WildR community include supplemented Brucella-blood agar [BBL ^TM^ Brucella broth (BD Cat#90000-076), 50 mg hemin/L, 0.5 mg menadione/L, and 5% defibrinated horse blood], Tryptone Yeast Extract Glucose (TYG) Medium (20g tryptone/L, 10g yeast extract/L, 5 g glucose/L, 1 g cysteine/L, 71 µM CaCl_2_, 78 µM MgSO_4_, 0.3 mM KH_2_PO_4_, 0.3 mM K_2_HPO_4_, 1.4 mM NaCl, 5 mg hemin/L, and 0.2% NaHCO_3_) (Bacic and Smith 2008), BHIS-FBS [Brain heart infusion (BHI) powder supplemented with 5 mg hemin/L, 0.2 mg menadione/L, 0.5 g cysteine/L, and 10% fetal bovine serum], BHIS-blood agar (BHI supplemented with 5 mg hemin/L, 1 g cysteine/L, 0.2% NaHCO_3_, 5% defibrinated horse blood, and 1.5% agar), Columbia-blood agar [Columbia broth powder (BD #294420) supplemented with 0.5 mg menadione/L and 5% defibrinated horse blood], MRS [MRS broth (BD #288130), supplemented with 1.5% agar if required], YCFAC [Yeast casitone fatty acids with carbohydrates broth (#AS-680) or agar (#AS-675) purchased from Anaerobe Systems], and BG [23.1 g Bryant and Burkey Medium (BB)/L (Sigma Aldrich #91903) and 12.5 g modified Gifu Anaerobic Medium (mGAM)/L (Himedia Laboratorires #M2079), which is a 70:30 ratio of BB:mGAM] (Javdan et al. 2020). All media was degassed for at least 24 hours before culturing.

Dilutions of the WildR reference stock or fresh fecal pellets collected from WildR-colonized mice resuspended in PBS were plated on anaerobic media and incubated anaerobically at 37 °C for 2-4 days. Single colonies were picked and used to inoculate broth cultures or were restruck on fresh plates and incubated at 37 °C anaerobically for 2 days. For storage, sterile, pre-reduced 50% glycerol was added to cultures, and the mixture was stored at-80 °C. Cultures were then screened for taxonomic identity by full-length 16S rRNA gene or *gyrB* gene sequencing (primer sequences provided in Supplemental Table 5. Target genes were amplified from ∼2 µL of cultured cells resuspended in 20 µL TE (10 mM Tris, pH 8; 1 mM EDTA). PCR products were cleaned up with Exo-SAP-IT (ThermoFisher, Catalog # 78201.1.ML) and then sequenced by Sanger sequencing. Resulting reads were trimmed and assigned putative taxonomy using blastn against the 16S ribosomal RNA sequences (Bacteria and Archea) database (Altschul et al. 1990). After initial screening, representative cultures from each species identified were subjected to an additional round of strain purification by streaking for single colonies, and species identity was reconfirmed by Sanger sequencing of the full-length 16S rRNA gene amplicons. Strains isolated and their growth conditions are listed in Supplemental Table 6.

### Whole genome sequencing of WildR isolates

De novo genome sequencing of WildR isolates was performed using a combination of short (Illumina) and long-read sequencing. High molecular weight gDNA was extracted from *Bacteroidales* sp. using the Qiagen genomic tip (20/G) kit following the standard protocol for Gram-negative extraction. The HMW DNA extraction kit from NEB (#T3060) was used for extracting HMW gDNA from *L. murinus* following the standard input protocol except the lysis buffer contained 5 µL of Qiagen lytic enzyme (catalog #158928) and 10 µL of metapolyzme (Millipore Sigma MAC4L-5MG resuspended in 750 µL of water). HMW gDNA was extracted from *L. reuteri* and *L. johnsonii* using the Promega Wizard HMW DNA Extraction kit (#A2920). For *M. intestinale*, *M. gordoncarteri*, *S. musculi*, and *D. freteri*, gDNA was extracted with mechanical lysis and column clean up. To lyse cells, 500 µL of PB (Qiagen), 250 µL 20% SDS, 550 µL of phenol:chloroform:isoamyl alcohol (24:24:1), and ∼250 µL of 0.1-mm-zirconia-silica beads were added to ∼2×10^9^ cells. Cells were then mechanically lysed with bead beating (RT, 1 min at high speed, 2 cycles) (BioSpec). Genomic DNA was further purified using columns from the Qiagen Blood and Tissue kit (#69504) and washed following the standard protocol. Extraction of genomic DNA from modified *B. acidifaciens* and *P. vulgatus* clones for resequencing was performed using the Qiagen Blood & Tissue kit (#69504) following the protocol for Gram-negative bacteria. For all gDNA extractions, the quality and size of gDNA was assessed by Qubit, nanodrop, and agarose gel electrophoresis before library prep.

Libraries for short-read whole genome sequencing were prepared with the Illumina DNA prep kit (catalog #20018704), multiplexed using barcoded adapters purchased from IDT (IDT for Illumina DN/RNA UD Indexes SetA, Catalog #20027213), and sequenced on either the Illumina iSeq or the MiniSeq, which included a 5% PhiX spike-in, to obtain paired-end reads (2×150 bp). To facilitate generation of closed genomes, *Bacteroidales* and *Lactobacillales* spp. were also sequenced on the Oxford Nanopore MinION. Libraries for long-read sequencing were prepared using LSK-110 with modifications to include ligation of barcodes (EXP-NBD104) following the protocol described for the LSK-109 kit. Briefly, DNA was repaired following the LSK-110 protocol using NEB buffers. Then, barcodes were ligated using Blunt/TA ligase following steps described for the LSK-109 kit. Finally, after barcode ligation, adapter sequences were ligated as described for LSK-110. The sequencing library was loaded onto a R.9.4.1 SpotON flow cell and run on the Mk1C until ≥ 200,000 reads were collected for each sample. If more reads were required, the flow cell was washed, and more libraries were loaded.

For *Bacteroidales* and *Lactobacillales* spp., genomes were assembled from long reads using Trycycler (v0.5.3, (Wick et al. 2021)). Briefly, reads were subsampled and assembled with either Flye (v2.9-b1768; (Kolmogorov et al. 2019)), Miniasm (v0.3-r179)+Minipolish (v0.1.2) (Wick and Holt 2019), or Raven (v1.8.1; https://github.com/lbcb-sci/raven) before combining similar assemblies to generate a single consensus sequence as described in the Trycycler wiki (https://github.com/rrwick/Trycycler/wiki). Assemblies were polished with Illumina reads using Polypolish (version 0.5.0; (Wick and Holt 2022)) and Pilon (version 1.24; (Walker et al. 2014)). For *Phocaeicola vulgatus*, genomes were assembled using the hybrid assembly feature in Unicycler (version 0.4.8; (Wick et al. 2017)). Finally, genomes were annotated with Prokka (version 1.14.6; (Seemann 2014)). For other species, adapters were removed from reads using BBDuk (version 38.84) in Geneious Prime (2024.0.5) (https://www.geneious.com), and then genomes were assembled with the Geneious assembler. For genomes deriving from modified clones of *B. acidifaciens* and *P. vulgatus,* spontaneous mutations were determined by aligning short reads to assembled genomes from the parental strains using BreSeq (Deatherage and Barrick 2014). Genome and assembly metrics for each isolate genome are listed in Supplemental Table 2, and all genomes have been deposited with links to BioProject PRJNA1367654 in the NCBI database.

### Determining the abundance of species in the WildR community

Abundance of species for which we had obtained a genome sequence (MAG or isolated-derived) was assessed by quantifying the frequency of metagenomic sequencing reads mapping to the genome of interest. Cleaned short, metagenomic reads from either wild donor, F2 WildR mice (Rosshart et al. 2017) or F7 WildR cryopreserved stocks were mapped to a catalog containing WildR F7 MAGs and isolate genomes using BWA-MEM (https://arxiv.org/abs/1303.3997), retaining only correctly paired read alignments using samtools (Li et al. 2009). Reads for all contigs in a MAG or isolate genome were summed and the percentage of reads per kilobase per million reads was calculated. For each sample, the total reads mapping to this catalog was retrieved using samtools flagstat. The same metagenomic reads were also mapped to all genomes in the CMMG catalog (https://ezmeta.unige.ch/CMMG/) using the same BWA-MEM command.

### Identification and bioinformatic characterization of an ICE-encoded T6SS in WildR genomes

To identify T6SS gene clusters present in WildR strains, we first searched for genes annotated to encode T6SS-related functions in the genomes of Bacteroidales isolates from the community. This led to the identification of candidate T6SS gene clusters in strains of *B. acidifaciens* and *B. caecimuris.* Blastp searches against the NCBI non-redundant protein sequence database with each of the candidate T6SS genes in the clusters found that each predicted T6SS component shares ≥80% identity with previously characterized T6SS proteins in other Bacteroidales species, and confirmed that the gene clusters contain the complete compliment of genes required to constitute a functional T6SS^iii^ (Altschul et al. 1990, Russell et al. 2014, Coyne et al. 2016). Putative effector genes were identified based on proximity of genes to the T6SS structural genes (e.g., VgrG) and the presence of N-terminal domains associated with T6SS effectors (e.g., PAAR). Effector activity of the C-terminal region of candidate effectors (∼100 aa) was predicted using CD-Search and Foldseek (Marchler-Bauer et al. 2009, van Kempen et al. 2023). For the later, positive hits from the AFDB50 database were defined by a probability score of 1. Immunity genes were assigned based on proximity to a putative effector, length (<200 a.a.), and presence of immunity domains identified by CD-Search.

The boundaries of the 115 kb T6SS-ICE were determined based on loss of 100% identity in a pairwise alignment between the *B. acidifaciens* and *B. caecimuris* F12 T6SS neighboring sequences (Geneious). Putative functions were assigned to each gene based on similar blastp hits (>30% identity) and classified according to functions typically encoded in ICEs (Coyne et al. 2014).

### *B. acidifaciens* methylome identification

For long-read sequencing using Pacific Biosciences (HiFi/SMRT sequencing), high-molecular-weight (HMW) genomic DNA from each isolate was extracted as described above for the Oxford Nanopore workflow. PacBio HiFi libraries were prepared using the SMRTbell Express Template Prep Kit v2.0 (Pacific Biosciences) according to the manufacturer’s recommendations for microbial HiFi genome sequencing, starting with 5–10 µg of HMW DNA per sample. Sequencing was performed on a PacBio Sequel II instrument at the Fred Hutchinson Cancer Center, generating circular consensus sequence (CCS) reads. Raw subreads were processed in SMRT Link (v11.0.0.146107), and CCS reads were used as input for the Microbial Assembly workflow, which performs long-read assembly, polishing, and contig finishing. For each genome, assembly statistics, CCS concordance, contiguity, depth of coverage, and base modification metrics were obtained directly from the SMRT Link analysis reports. Methylation analysis was carried out using the Base Modification and Motif Analysis module with default m6A/m4C detection models. Final polished assemblies and methylome calls for all strains (*B. acidifaciens*, *B. caecimuris* F12, *B. thetaiotaomicron*, *B. uniformis*, *P. vulgatus*, and *P. distasonis*) were exported from SMRT Link and used for downstream comparative analysis. Genome assemblies have been deposited with links to BioProject PRJNA1367654 in the NCBI database.

### Genetic manipulation of *Bacteroides* strains

Standard molecular procedures were employed for creation, maintenance and transformation of plasmids in *E. coli.* The primers, plasmids, and strains designed for this study are listed in Supplemental Table 5. All primers were purchased from Integrated DNA Technologies (IDT) and geneblocks were synthesized by Twist Biosciences. Enzymes such as OneTaq polymerase, restriction enzymes, and Gibson Assembly reagents were purchased from New England Biolabs (NEB). To generate pNBU2-erm-us1311::*tssC*-singleRMsilent, a two bp change was introduced in the GATATC motif found in the plasmid using quickchange. To generate pLGB13-RMsilent and pNBU2-erm-RMsilent::P5E4, all *B. acidifaciens* methylation sites were identified in the parent plasmid by sequence homology (eight and four, respectively) and then single base pair mutations were designed to remove specific motifs. Plasmids were assembled from synthesized geneblocks containing mutations and amplified fragments of unmodified regions using Gibson assembly and validated by whole plasmid sequencing (Plasmidsauraus).

Gene knockouts in *B. acidifaciens* using either pSIE1 or pLGB13 derivatives were generated following previously published protocols with the following modifications (Garcia-Bayona and Comstock 2019, Bencivenga-Barry et al. 2020). Briefly, plasmids were mobilized from *E. coli* S17 into *B. acidifaciens* via overnight anaerobic mating at 37 °C where *E. coli* was mixed with a 10-fold excess of *B. acidifaciens*. To further improve mating efficiency, *E. coli* S17 harboring the RK231 helper plasmid was also included in the mating mixture at equal CFU to *E. coli* S17 harboring the plasmid for allelic exchange (Smith et al. 1992). Merodiploids were isolated by plating on selective media, passaged once in BHIS-FBS without antibiotics to allow for plasmid excision, and plated on Brucella-blood supplemented with aTc for counter selection of the plasmid. Resulting colonies were patched onto selective Brucella-blood plates to confirm loss of the plasmid, and the gene disruption of interest was confirmed by colony PCR. For select mutants employed in germ-free mouse colonization experiments, we performed whole-genome sequencing using an Illumina MiniSeq to identify second site mutations, as described above. Strains expressing heterologous copies of *clpV* were generated by cloning *clpV* and an intrinsic terminator from pWW3855 downstream of the P5E4 promoter in pNBU2-ermG-RMsilent::P5E4 as previously described (Whitaker et al. 2017).

Integration of pNBU2-ermG or derivatives was performed as previously described (Wang et al. 2000, Koropatkin et al. 2008, Whitaker et al. 2017), but with modified mating conditions as indicated above. Resulting transconjugants were selected on Brucella-blood supplemented with gentamycin and erythromycin, colony purified to confirm Erm^R^, and stocked.

### Interbacterial transfer of the GA1 T6SS-encoding ICE

To transfer the T6SS-encoding ICE from *B. acidifaciens* to *P. vulgatus*, we modified previously described methods for spontaneous *in vitro* mobile element transfer (Supplemental Figure 4B; (Ross et al. 2019)). First, we used allelic exchange as described above to insert a chloramphenicol resistance gene under the control of the constitutive P1 promoter in a neutral site in the ICE in *B. acidifaciens*. Then, Cm^R^ *B. acidifaciens* cells from overnight cultures were mixed with Erm^R^ *P. vulgatus* cells at a 10:1 ratio (OD_600_ 6) on Brucella-blood plates. Following 24-hours of anaerobic co-culture growth, cells were harvested, resuspended in PBS, and plated on Brucella-blood supplemented with chloramphenicol and erythromycin to select transconjugants. Resulting transconjugants were colony purified and whole genome sequencing was performed using the Illumina platform as described above. Resulting reads that aligned to both the ICE and *P. vulgatus* genome, determined using minimap2 were used to identify the ICE insertion sites (Li 2018). Reads generated in this sequencing have been deposited with links to BioProject PRJNA1367654 in the NCBI database.

### *In vitro* bacterial growth competition assays

For intra-and interspecies growth competition experiments, donor and recipient strains were spread as lawns on plates and incubated anaerobically at 37 °C for 24-48 hours. Most *Bacteroides* sp, *Parabacteroides distasonis*, and *Phocaeicola vulgatus* were grown on Brucella-blood plates, except for *B. sp*910578895, which was grown on BHIS-blood. Other media such as Columbia-agar (*S. musculi*, *M. intestinale*), MRS agar (*L. murinus*), LB agar (*E. fergusonii*), and YCFAC (*D. freteri*, *M. gordoncarteri*) were used to culture additional recipient species tested (Supplemental Table 6). Cells were harvested from plates and resuspended in 1 mL PBS, before measuring the OD_600_ and diluting to OD 20 or OD 0.2 for donor and recipient strains, respectively. Cell dilutions were combined at a 1:1 ratio (100:1 final donor:recipient ratio), 10 µL of the mixture was spotted onto plates, and mixtures were incubated anaerobically at 37 °C for 16-20 hours. Most assays were performed on Brucella-blood plates supplemented with 3% agar except for competitions performed on BHIS-blood with 3% agar (*B*. *sp*910578895 and *E. gallinarum*), Columbia-blood with 3% agar (*S. musculi* and *M. intestinale*), or YCFAC (*M. gordoncarteri*, *D. freteri*). Growth competitions with strict anaerobes (*S. musculi*, *M intestinale*, *M. gordoncateri*, and *D. freteri*) were performed entirely under anaerobic conditions using pre-reduced media. After incubation, all cells were harvested and resuspended in 1 mL of PBS. Quantification of species abundance at the start and end of each assay was performed by enumerating colony forming units (CFU) or qPCR. CFU enumeration was used to quantify species abundance for intraspecies (recipients, Em^R^) or interspecies (donors, *B. acidifaciens* Cm^R^; recipients *P. vulgatus*, *B. uniformis*, *B. thetaiotaomicron*, *B. caecimuris* F5, *B. caecimuris* F12, and *P. distasonis*, Em^R^ or *E. gallinarum* (BHI-blood, aerobic), *L. murinus* (MRS, anaerobic), or *E. fergusonii* (LB, aerobic)) growth competitions. Interspecies growth competitions with strict anaerobes, *B. sp*910578895, or *P. vulgatus* donors were quantified by qPCR using strain specific primers (Supplemental Table 5). For qPCR, genomic DNA from cell pellets of competition mixtures was extracted using either the Qiagen Blood & Tissue kit following the protocol for extraction from Gram-negative organisms or via mechanical lysis as described above. DNA was quantified via Qubit (ThermoFisher) and diluted before using as template DNA in qPCR (SsoAdvanced, BioRad). All experiments were performed with technical triplicates for at least three biological replicates.

### Gnotobiotic mouse colonization studies

#### Animal care

In accordance with protocols approved by the University of Washington Institutional Animal Care and Use Committee, germ-free 5-12 week old male and female Swiss Webster mice from multiple litters were randomly selected and housed in pairs in single Techniplast cages with a 12-hour light/dark cycle and fed a standard lab diet (Laboratory Autoclavable Rodent Diet 5010, LabDiet). Blinding was not performed. For each experiment, the number of animals per group (n=4) was selected based on Lemorte power calculations to measure a difference of ∼10-fold bacterial abundance with a *p* < 0.05 at 80% power along with housing and maintenance considerations for gnotobiotic mice. Mouse feces were confirmed to be sterile prior to colonization by PCR with primers targeting the 16S rRNA gene.

#### Mouse colonization procedures

Bacterial strains to be employed in germ-free mouse colonization experiments were grown anaerobically on Brucella-blood plates, resuspended in BHI broth, and diluted. For experiments to determine strain competitiveness in a simplified context, pairs of strains were combined at a 100:1 (10^9^:10^7^ CFU, *B. acidifaciens: P. vulgatus*) or 10:1 ratio (10^9^: 10^8^ CFU, all other donor–recipient pairs), as determined by prior calculation of the OD_600_ to CFU correspondence. This mixture was introduced into germ-free mice via oral gavage (200 µL per mouse).

To colonize mice with the WildR microbiome and a genetically modified WildR isolate (e.g., *B. acid ^exo^*), we first estimated the approximate abundance of the endogenous isolate (e.g., *B. acid ^endo^)* in the WildR stocks using a combination of CFU enumeration, qPCR, and 16S rRNA sequencing. The total CFU of Bacteroidales strains from the *Bacteroides*, *Phocaeicola*, and *Parabacteroides* in the WildR reference stocks was determined by plating on Brucella-blood agar supplemented with gentamycin, a condition selective for the order. The abundance of *B. acidifaciens* relative to total Bacteroidales levels in WildR stocks was then determined using qPCR with species and order-specific primers on DNA extracted from WildR reference stocks, and this measurement was used to estimate the contribution of CFUs from *B. acidifaciens* to the total Bacteroidales population (∼3×10^4^ CFU/ml of WildR cryopreserved stocks). An estimate for the abundance of *P. vulgatus* CFUs in WildR stocks (∼3×10^6^ CFU/ml) was calculated in a similar manner, except the fraction of total Bacteroidales represented by this strain was determined by counting reads from 16S rRNA gene sequencing of amplicons from WildR reference stock genomic DNA, as described below.

Modified strains to be introduced to mice along with the WildR community were grown and suspended in BHI broth as described above for isolate colonization studies. Suspensions were then diluted to the desired CFU level (∼3×10^4^, 3×10^5^ or 3×10^6^ *B. acid ^exo^* CFU or ∼3×10^7^ *P. vul ^exo^* CFU) in 1 mL of the WildR reference stock under anaerobic conditions. After preparation, the mixture was stored in Wheaton crimp top vials to maintain an anaerobic environment and then introduced into germ-free Swiss Webster mice via a single oral gavage (150-200 µL per mouse). For all germ-free mouse colonization experiments, levels of introduced strains in the gavage mixtures were determined by diluting and plating on selective media.

#### Quantification of bacterial populations in fecal and cecal samples

Fresh fecal pellets were collected from colonized animals at regular intervals, and cecal contents were collected from euthanized animals at the conclusion of the experiment as described for WildR reference stock generation. To enumerate CFUs of marked strains, fresh fecal pellets or cecal contents (∼50-100 mg/sample) were homogenized in PBS using sterile pestles. Insoluble material was removed via gentle centrifugation (500 x*g*, 30s) and the resulting supernatant was diluted in PBS. Dilutions were plated on selective media (e.g., Brucella-blood supplemented with erythromycin and gentamycin to isolate *B. acid ^exo^* from the WildR community) and incubated anaerobically at 37 °C. Final CFU counts were normalized to initial fecal weights. Remaining fecal or cecal material from these collections was stored at-80 °C for gDNA extraction.

The abundance of modified strains relative to total bacteria present or endogenous populations in WildR-colonized mice was determined by qPCR. Genomic DNA was extracted from thawed fecal pellets or cecal contents (∼50-100 mg/sample) by first combining samples with 200 µL of 0.1-mm-diameter zirconia/silica beads (BioSpec Products, Bartlesville, OK), 500 µL of CP Buffer (Omega Biotek, #PDR042) or PB Buffer (Qiagen), 250 µL of 20% SDS, and 550 µL of phenol:chloroform:isoamyl alcohol (24:24:1) (ThermoFisher). Microbial cells were mechanically lysed with a bead beater at room temperature (BioSpec Products; instrument on high for 1 min; 2 cycles) and then centrifuged at 10,000 *xg* for 10 min to separate phases. All of the aqueous phase containing gDNA (∼400-500 µL/sample) was applied to E-Z 96^TM^ DNA plates (Omega Biotek; #BD96-01) or Qiagen Blood and Tissue columns and cleaned following the standard protocol. DNA was quantified by fluorimetry (Qubit, ThermoFisher) and then diluted to ∼5 ng/uL in EB (10 mM Tris-HCl pH 8.5; Omega Biotek PDR048). Relative species or strain abundance was determined by qPCR using primers described in Supplemental Table 5. All qPCR was performed using a CFX96 instrument (BioRad) and SsoAdvanced Universal SYBR Green Supermix (BioRad).

#### Community structure analysis by 16S rRNA gene amplicon

The V3-V4 region of the 16S rRNA gene was amplified from genomic DNA extracted from fecal and cecal samples following the standard Illumina protocol for 16S metagenomic sequencing (Illumina Part # 15044223 Rev. B) with the following modifications. Briefly, purified gDNA was quantified by Qubit, diluted to 5 ng/µL, and amplified using primers 16S_v3v4_fwd_Illumina and 16S_v3v4_rev_Illumina (Supplemental Table 5) and 2x KAPA HiFi HotStart ReadyMix (Roche). PCR was performed in triplicate to avoid jackpot amplification. Reactions were prepared in a biosafety cabinet to prevent contamination and each run included no-template controls. Pooled PCR products for each sample were cleaned using AmpureXP beads (Bruker). A second PCR was performed to add sequencing adapters and barcodes (IDT for Illumina UD Indexes Plates A-D) using 2x KAPA HiFi HotStart ReadyMix. Final PCR products were cleaned and normalized with the Sequalprep Normalization Plate kit (Thermo Scientific). The size and quantity of DNA libraries from select samples was confirmed by agarose gel electrophoresis and fluorimetry (Qubit), respectively.

Equal volumes of 16S rRNA libraries prepared from up to 384 samples was pooled and quantified for sequencing. Libraries were denatured, diluted, and combined with PhiX spike-in before sequencing on the MiSeq or NextSeq2000 (300-bp paired reads; dual 10 bp indexes; 10 or 25% PhiX, respectively). Samples were demultiplexed with onboard software for each instrument. All data generated from this sequencing has been deposited with links to BioProject PRJNA1367654 in the NCBI database.

Analysis of 16S rRNA sequencing data was performed using Qiime2 (v2020.11; (Bolyen et al. 2019)). Primers were trimmed using the qiime cutadapt plugin before using DADA2 for ASV calling and counting to determine the relative abundance of each ASV. Beta diversity metrics (Unifrac) were calculated from rarefied data using the qiime diversity plugin and statistical tests were performed with the qiime longitudinal plugin. Principle component analysis plots were generated in R using cmdscale. To quantify total *B. acidifaciens* abundance, we performed a custom taxonomic assignment of ASVs. The top taxonomic hit for each ASV was obtained by a BLAST search of each ASV in the WildR MAG and isolate genome catalog. These new taxonomic assignments were combined with relative abundance of each ASV calculated from the DADA2 analysis to determine relative frequencies of *B. acidifaciens*.

### ICE-seq library preparation and data analysis

Genomic DNA was extracted from fecal samples as described above and used to prepare ICE-seq libraries following previously described methods with the modifications described below (Gallagher 2019) (Supplemental Figure 4A). Briefly, DNA was sheared to ∼500 bp average size and C-tailed. Sequences outside of the ICE at either the 5’ or 3’ ends of the ICE were specifically amplified using primers annealing within the ICE at either the 5’ or 3’ end (“ICE junction”) and a C-tailed specific primer (Supplemental Table 5). After DNA purification, a second round of PCR was performed to amplify the ICE junctions and add adapter and index sequences for sequencing. Samples were pooled and multiplexed samples were sequenced as 75-bp single-end reads on an Illumina MiniSeq with 30% PhiX DNA spiked in using a custom mix containing three primers for sequencing the 5’, 3’, and PhiX sequences.

Illumina sequencing reads were analyzed using a portion of previously described custom Python script (Gallagher 2019). Sequences were filtered for those containing the first 5 bp (ATAAT) or 7 bp (ATTATTG) of the ICE 5’ and 3’ ends, respectively. Then reads were aligned using a BWA aligner to a composite CMMG-WildR catalog, which was dereplicated (at 99% ANI) using dRep (v3.5.0) (Olm et al. 2017, Kieser et al. 2022). Reads per insertion site were tallied, and insertion sites were filtered to include only sites where reads mapped in opposite directions from both ends of the ICE. To calculate the percentage of ICE reads mapping in a sample, the total of mapped reads fitting these criteria was calculated and the fraction of each insertion site was then determined. The percentage was independently calculated for reads generated from either end of the ICE. Sequencing reads have been deposited with links to BioProject PRJNA1367654 in the NCBI database.

## Statistical analysis

Statistical analysis was performed using GraphPad Prism (version 10.2.3 for Mac OS X, GraphPad Software, Boston, Massachusetts USA, www.graphpad.com) using tests described in the main text and figure legends.

---

**Supplemental Figure 1.**
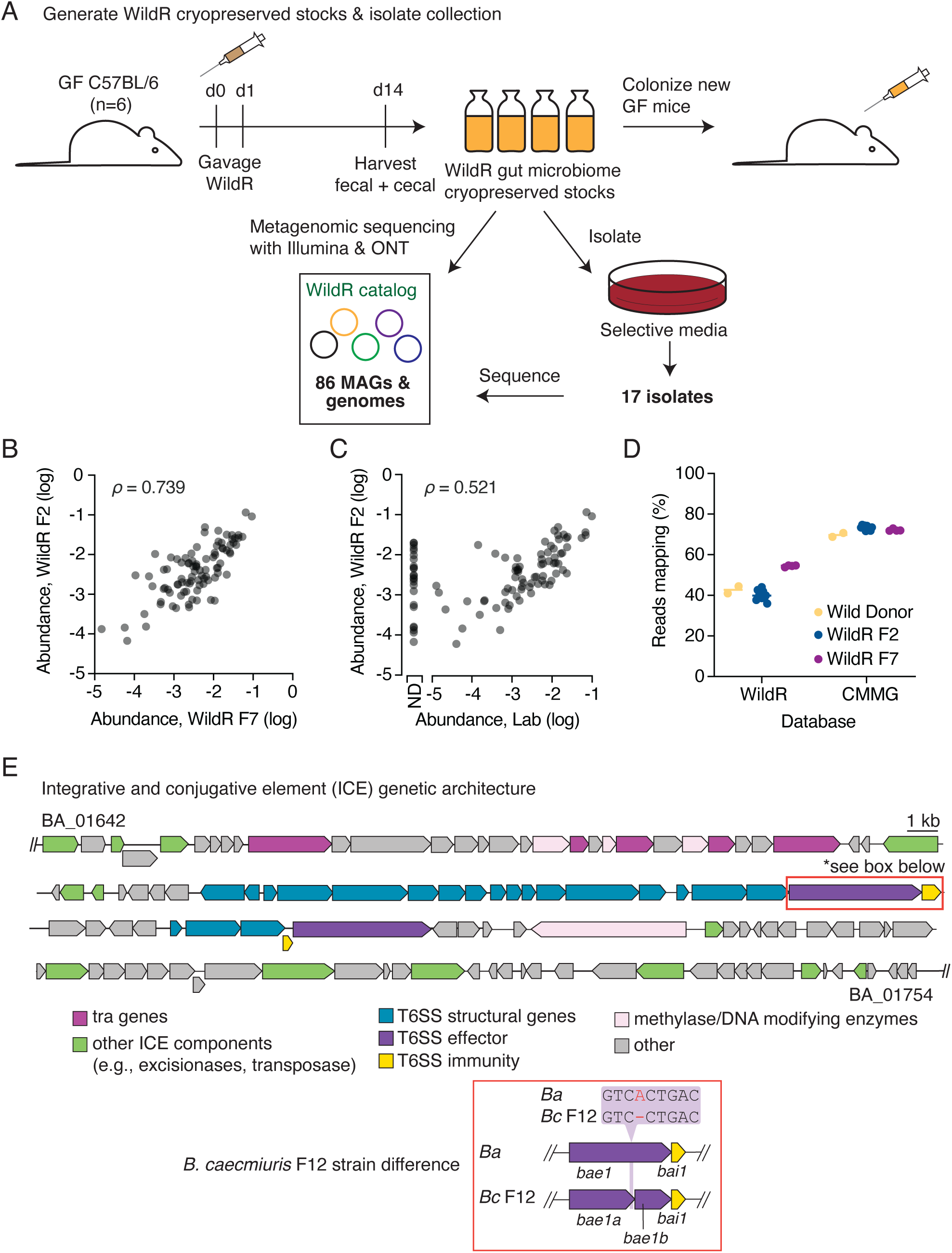
E**v**idence **supporting stability of the WildR microbiome over multiple generations and identification of an ICE-encoded T6SS** (A) Schematic depicting the generation of the WildR F7 cryoprserved stocks from the combined cecal contents of six mice used to propagate the community, and subsequent community characterization steps. (B-C) Comparison of the abundance of the 100 genera most prevalent in wild donor mice between the indicated communities. Points are shaded to show overlapping datapoints (darker shades), and Spearman’s correlation for each comparison is indicated (π). ND, not detected. (D) Mapping efficiency of metagenomic reads from different WildR community generations to selected murine-derived genome databases: the WildR catalog (86 MAGs and genomes from the WildR, generated in this study) or the comprehensive mouse microbiota genome catalog (CMMG; 1573 species across mouse microbiomes (Kieser et al. 2022)). (E) To-scale schematic of the ICE containing the GA1 T6SS encoded in *B. acidifaciens* and *B. caecimuris* F12. Location of single base deletion in *B. caecimuris* F12 highlighted in the orange box.

**Supplemental Figure 2.**
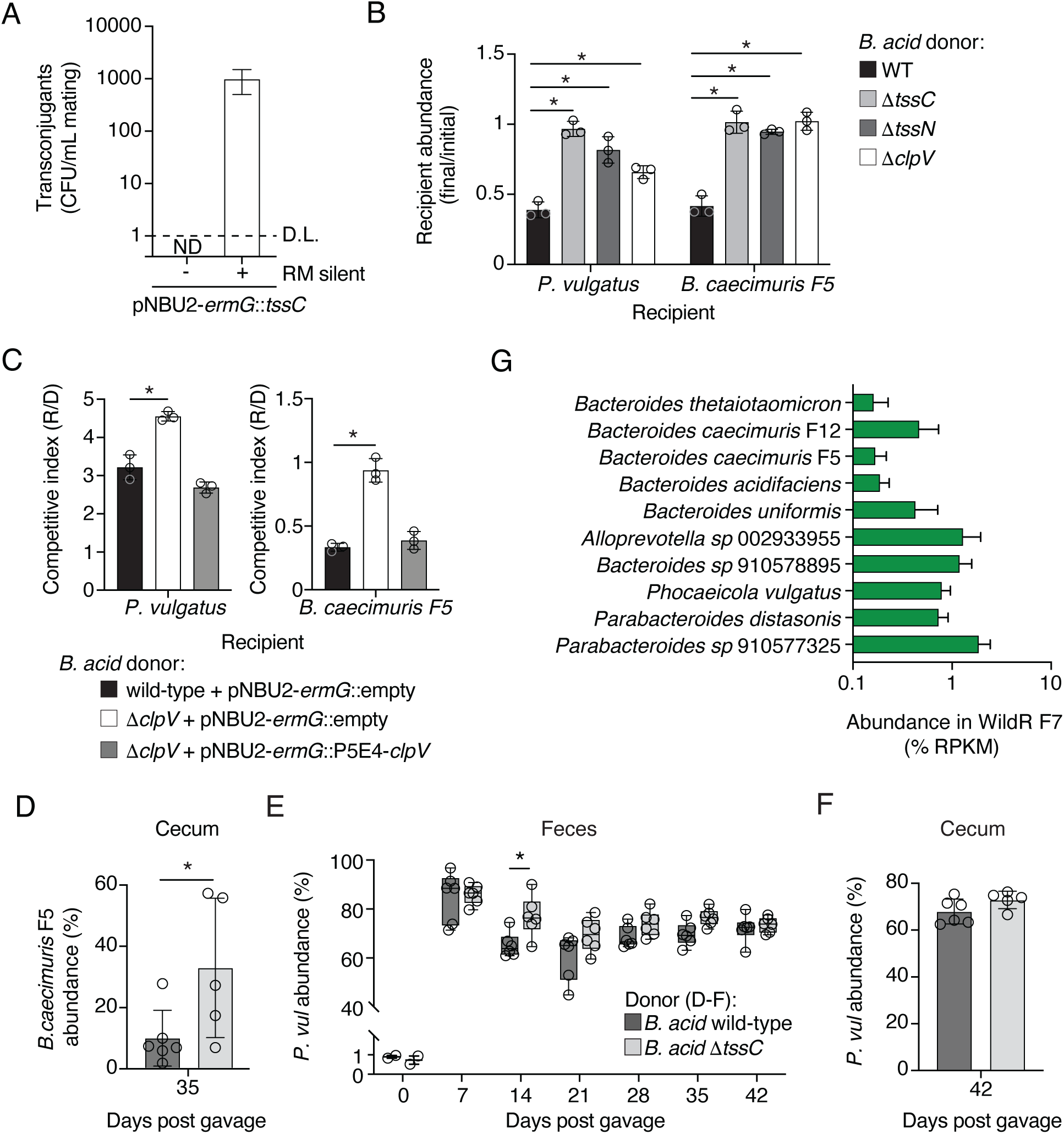
A**c**tivity **of the *B. acidifaciens* T6SS against species co-resident in the WildR community.** (A) *In vitro* mating efficiency of the integrative plasmid pNBU2-*ermG:: tssC* into *B. acidifaciens*. “RM silent” indicates plasmid was mutated to remove a *B. acidifaciens* methylated motif. Data shown are mean ± SD from 3 independent matings. N.D., not detected; D.L., detection limit. (B) Recipient abundance after *in vitro* growth competitions between *B. acidifaciens* donors lacking various structural components of the T6SS and the indicated recipient species. (C) Competitive index from *in vitro* growth competition between indicated WildR isolates (recipient) and *B. acidifaciens* donors. For B & C, data show the mean ± SD of technical replicates from one biological replicate and represent results from at least three biological replicates. \**p* ≤ 0.05 by two-tailed t-test; all other comparisons were not significant. (D) *B. caecimuris* F5 abundance in cecal contents from germ-free mice co-colonized with *B. acidifaciens* wild-type or Δ*tssC*. Data show the mean ± SD and points indicate values from individual mice (n=6) across two biological replicates. \**p* ≤ 0.05 (two-tailed t-test). (E) *P. vulguatus* abundance in feces from germ-free mice (n=6, two biological replicates) co-colonized with *B. acidifaciens* wild-type or Δ*tssC* and *P. vulguatus*. Boxplots represent the interquartile range with indicated mean for each condition, whiskers represent minimum and maximum values, points represent individual values. \**p* ≤ 0.05 (two-way ANOVA with repeated measures test and Šidák’s multiple comparisons test). (F) *P. vulguatus* abundance in cecal contents from mice described in panel E. Data show the mean ± SD and points show values from individual mice (n=6). No statistical difference was found based on *B. acidifaciens* genotype by two-tailed t-test. (G) Relative abundance (% reads per kb per million) of selected WildR Bacteroidales species in the WildR F7 generation. Data are mean + SD from cryopreserved WildR stocks and 2 fecal samples from mice used to propagate the WildR F7 community.

**Supplemental Figure 3.**
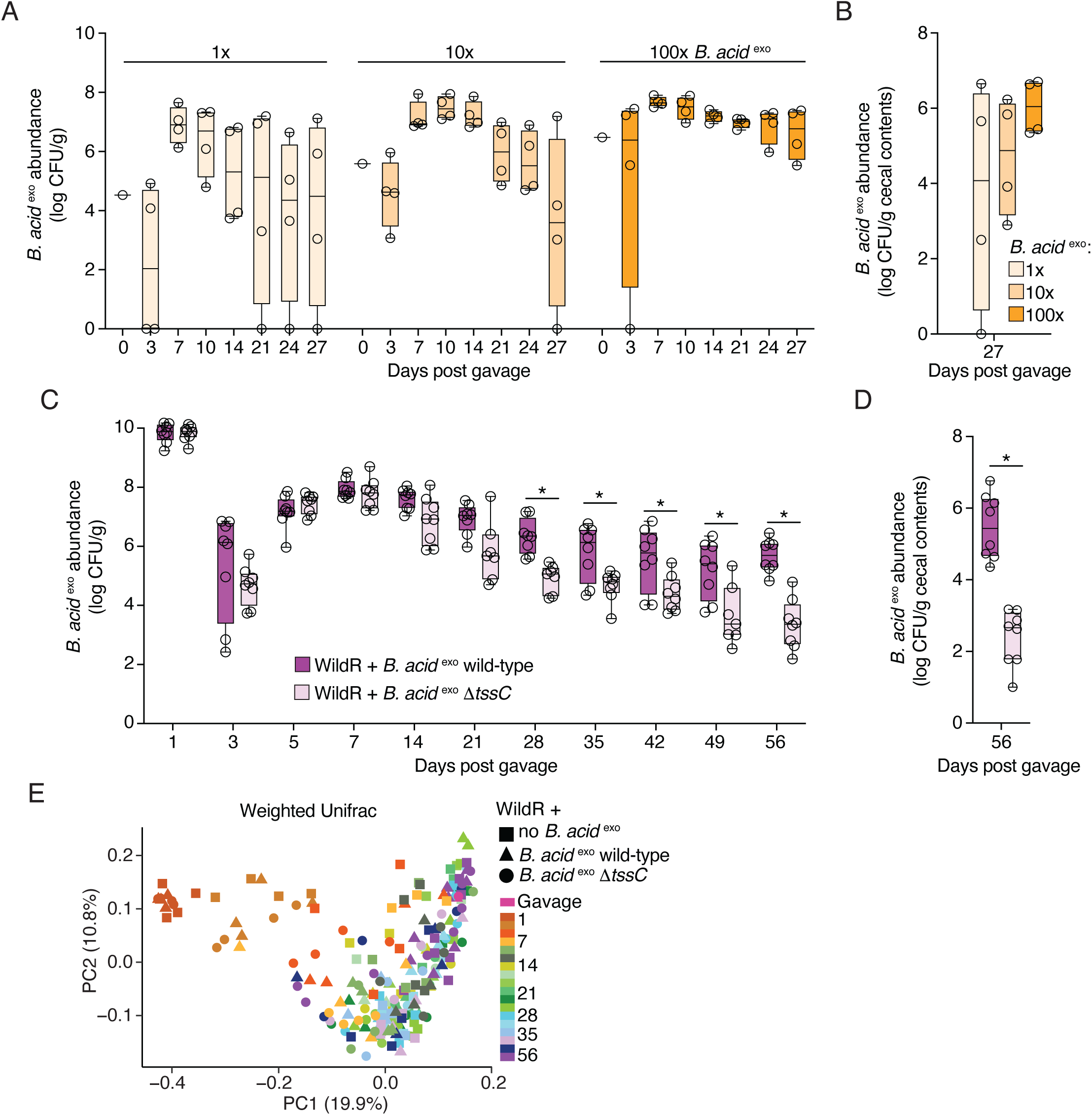
Addition of *B. acid* ^exo^ to the WildR does not alter community composition and enables *in situ* measurement of T6SS-mediated fitness. (A,B) Recovery of *B. acid* ^exo^ from feces (A) and cecal contents (B) following gavage of germ-free mice with the WildR community and the indicated amount of *B. acid* ^exo^ relative to the endogenous population. (C,D) Recovery of *B. acid* ^exo^ or *B. acid* ^exo^ Δ*tssC* from feces (D) or cecal contents (E) from mice colonized with the WildR and *B. acid* ^exo^ strains. \**p* ≤ 0.05 (two-way ANOVA with repeated measures test and Šidák’s multiple comparisons test in C, unpaired t-test in D). For data in panels A-D, boxplots represent the interquartile range with indicated mean for each condition, whiskers represent minimum and maximum values, and points show values from individual mice. (E) Principal coordinate analysis of weighted Unifrac diversity metrics calculated from 16S rRNA amplicon sequencing data from feces collected from mice colonized with the WildR alone or in combination with the indicated strain of *B. acid* ^exo^. Gavage samples highlighted in pink and remaining timed fecal and cecal (collected at 56 days post gavage) samples are colored as indicated. Data shown are from one biological replicate, representative two experiments conducted.

**Supplemental Figure 4.**
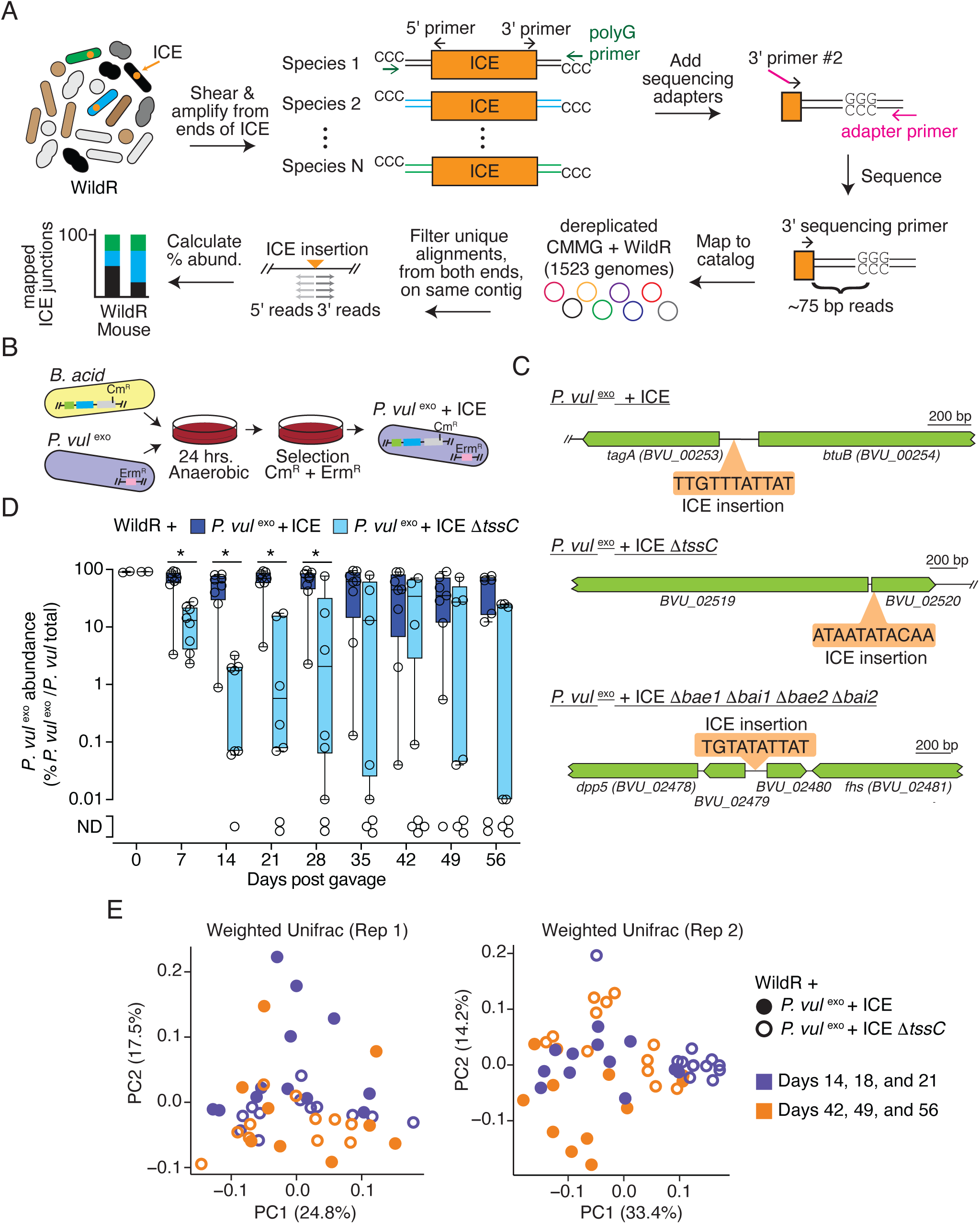
D**i**stribution **of the T6SS-ICE in the WildR suggests limited fitness benefit to some *Bacteroides* sp. (**A) Schematic of ICE-seq approach to identify WildR species encoding the ICE. The junction amplification and sequencing strategy applied to both ends of the ICE is depicted only for the 3′ end for simplicity. (B) Schematic depicting ICE transfer from *B. acidifaciens* (marked with Cm^R^) to *P. vulgatus* (marked with Erm^R^) via *in vitro* conjugation and selective plating. (C) To-scale schematic of ICE insertion sites in *P. vulgatus* ^exo^ + ICE transconjugants that acquired the indicated versions of the ICE. (D) Abundance (relative to total *P. vulgatus*) of the indicated *P. vulgatus ^exo^* in fecal samples from mice co-colonized with the WildR community, as determined by qPCR. Boxplots represent the interquartile range with indicated mean for each condition, whiskers represent minimum and maximum values, and points show values from individual mice (n=8) from two biological replicates. Asterisks indicate significant differences between *P. vul* ^exo^ + ICE and *P. vul* ^exo^ + ICE Δ*tssC* frequency at the indicated time points (*p*<0.05, Šídák’s multiple comparisons test, mixed model ANOVA with multiple comparisons). N.D., not detected. (E) Principal coordinate analysis of weighted Unifrac diversity metrics calculated from 16S rRNA gene amplicon sequencing data from feces collected from mice (n=8/group across 2 biological replicates) colonized with the WildR and either *P. vul* ^exo^ + ICE (closed circles) or *P. vul* ^exo^ + ICE Δ*tssC* (open circles). The community composition varied between groups at early time points (purple; Rep 1, *p*=0.021, pseudo-F=2.7; Rep 2, *p*=0.011, pseudo-F=12, PERMANOVA test), but varied less or not signifcantly at late time points (orange; Rep 1, *p*=0.25, pseudo-F=1.3; Rep 2, *p*=0.003, pseudo-F=5).

